# Induced RPB1 depletion reveals a direct gene-specific control of RNA Polymerase III function by RNA Polymerase II

**DOI:** 10.1101/764605

**Authors:** Alan Gerber, Keiichi Ito, Chi-Shuen Chu, Robert G. Roeder

## Abstract

Increasing evidence suggests that tRNA levels are dynamically and specifically regulated in response to internal and external cues to modulate the cellular translational program. However, the molecular players and the mechanisms regulating the gene-specific expression of tRNAs are still unknown. Using an inducible auxin-degron system to rapidly deplete RPB1 (the largest subunit of RNA Pol II) in living cells, we identified Pol II as a direct gene-specific regulator of tRNA transcription. Our data suggest that Pol II transcription robustly interferes with Pol III function at specific tRNA genes. This activity was further found to be essential for MAF1-mediated repression of a large set of tRNA genes during serum starvation, indicating that repression of tRNA genes by Pol II is dynamically regulated. Hence, Pol II plays a direct and central role in the gene-specific regulation of tRNA expression.

## Introduction

Eukaryotic nuclear transcription is carried out by RNA polymerases I, II and III (Pols I, II and III). Pol I transcribes the single multi-copy gene that specifies the large ribosomal RNAs (28S rRNA, 18S rRNA, and 5.8S rRNA). Pol II transcribes genes that specify pre-messenger RNAs and long non-coding RNAs (the vast majority), as well as various small functional RNAs such as microRNAs and small nuclear RNAs (snRNAs). Finally, Pol III transcribes genes that specify small untranslated RNAs essential for protein synthesis, such as transfer RNAs (tRNAs) and the 5S ribosomal RNA, as well as a heteroclite group of short regulatory RNAs (Palazzo and Lee, 2015; Roeder and Rutter, 1969). In contrast to transcription by Pol II, which involves countless types of promoters and enhancers and many diverse transcriptional regulators, transcription of Pol III genes involves only three types of promoters that are regulated by a limited set of transcription factors (Dumay-Odelot et al., 2014; Schramm and Hernandez, 2002). Notably, transcription of all tRNA genes, other than those for selenocysteine tRNAs, is governed by type II internal promoters that require only the conserved multi-subunit transcription factors TFIIIB and TFIIIC in conjunction with Pol III. Initiation of tRNA transcription starts with the binding of TFIIIC to two intragenic boxes in the tRNA coding sequence. TFIIIC, in turns, promotes the association of TFIIIB to a region immediately upstream of the TSS. Finally, TFIIIB recruits Pol III to direct the first round of transcription. Upon reaching the terminator, a stretch of 4 or more thymidine residues, Pol III can be recycled to the same TFIIIB-bound promoter at much higher rates than the initial preinitiation complex (PIC) formation, in a process known as facilitated recycling (Arimbasseri et al., 2014; Orioli et al., 2012).

tRNAs are stable molecules synthesized at very high rates (tens of millions of tRNAs every cell cycle in mammalian cells). Their expression is co-regulated, primarily at the level of transcription initiation, in response to nutrient availability and cellular stresses via different signaling pathways (Grewal, 2015; Moir and Willis, 2013). Despite the apparent simplicity in the Pol III transcriptional machinery, the relative abundance of individual tRNAs varies considerably across different tissues and cell lines (Dittmar et al., 2006; Sagi et al., 2016) in response to stresses and changes in growth conditions (Cieśla et al., 2007; Orioli et al., 2016; Pang et al., 2014; Torrent et al., 2018), in support of cell proliferation or differentiation (Gingold et al., 2014) and in pathologies such as cancer (Goodarzi et al., 2016; Pavon-Eternod et al., 2009). Dynamic changes in the composition of the cellular tRNA repertoire in response to external and internal cues may in turn affect protein abundance, translation fidelity, protein folding and even mRNA stability (reviewed in (Rak et al., 2018) to modify cellular metabolism or physiology.

The levels of a specific tRNA are determined by both its stability and its synthesis rate (Rak et al., 2018; Wichtowska et al., 2013). Surprisingly, however, there are no known tissue- or gene-specific transcription factors dedicated to tRNA gene transcription. Interestingly, in human and mouse cells, Pol II is often found in close proximity to tRNA genes that display high Pol III occupancy (Barski et al., 2010; Canella et al., 2012; Moqtaderi et al., 2010; Oler et al., 2010; Raha et al., 2010; Yeganeh et al., 2017). Different hypotheses have been proposed to explain these observations, and range from a spurious recruitment of Pol II at genes displaying high levels of Pol III (Canella et al., 2012; Yeganeh et al., 2017) to a requirement of Pol II activity to remodel the chromatin environment in order to facilitate Pol III recruitment or activity. Treatment of cells with the transcriptional inhibitor α-amanitin at a concentration selective for Pol II was found to mildly reduce the expression of some Pol III targets (Barski et al., 2010; Listerman et al., 2007; Raha et al., 2010). However, and especially in light of the slow action of α-amanitin in vivo (Nguyen et al., 1996), these experiments did not establish that these effects were direct. In contrast, the expression levels of the Pol II-transcribed *AtNUDT22* gene in Arabidopsis negatively correlate with the expression of a set of overlapping Pol III-transcribed proline tRNA genes (Lukoszek et al., 2013). These results suggest transcriptional interference between the two types of polymerases rather than Pol II-assisted enhancement of Pol III activity. In order to unravel the primary function of Pol II at Pol III-transcribed genes *in vivo*, we established a human cell line that allows an inducible, rapid and selective depletion of RPB1 in living cells using an auxin-degron system.

Here, by exploiting this auxin-inducible system along with different Pol II transcriptional inhibitors, we report the identification of specific tRNA genes whose transcription by Pol III is directly under the control of elongating Pol II complexes. Unexpectedly, we found that the auxin-induced degradation of RPB1 in Pol II complexes was incomplete, generating termination incompetent CTD-less (Pol IIB) complexes specifically at small nuclear RNA (snRNAs) and stably paused genes. By comparing the changes in the levels of nascent tRNAs in cells treated with the pause-release inhibitor DRB and in cells after auxin-induced RPB1 degradation, we further confirmed that Pol II passage through tRNA genes interferes robustly with their transcription. Pol II-mediated repression of Pol III also appeared to be essential for MAF1-mediated repression of a majority of tRNA genes during serum starvation. In addition, our experiments reveal that the downregulation of specific tRNA genes following Pol II inhibition or depletion is caused mainly by indirect effects such as loss of the MYC protein, which can explain the reported downregulation of particular tRNA genes in α-amanitin-treated cells (Barski et al., 2010; Raha et al., 2010). Hence, Pol II can regulate tRNA transcription by Pol III dynamically and in a gene-specific manner both directly, through transcriptional interference, and indirectly, by regulating the expression of MYC.

## Results

### Establishment of a cell line for rapid and inducible depletion of RPB1

To study the interplay between Pol II and Pol III, we established a HEK293 cell line expressing a full length RPB1 that is C-terminally tagged with a minimal auxin-sensitive degron (Natsume et al., 2016), and an orange fluorescent protein (**Figure 1A**) as the sole source of this essential subunit of Pol II as described in **Figure S1A-C**. In these cells, a rapid and auxin-dependent degradation of RPB1 was observed when the expression of the plant ubiquitin ligase OsTIR1 was induced for 12 hrs with doxycycline (Dox) followed by addition of auxin for 2 hrs (**Figure 1B**). Three clones (#7, #12 and #19) displaying high responsiveness to the treatment were selected and used in all further experiments (**Figure 1C**). A temporal analysis of these clones by flow cytometry revealed a rapid decay of fluorescence with a half-life of 33.2 min ± 1.8 min following auxin addition (**Figure 1D-F**). Microarray analysis of pools of RNA isolated from all three clones in untreated (Ctl), DOX-treated (D) or DOX treatment followed by auxin (DA) conditions revealed that steady-state levels of Pol II and Pol III transcripts were largely unchanged (**Figure 1G-I**). Surprisingly, about two third of the genes that were affected more than 2-fold (205/331 genes) were even upregulated in RPB1-depleted cells (**Figure 1I**). Pol II depletion on target genes was confirmed by RPB3 ChIP-seq for clones #7 and #19, which are, respectively, the most and least auxin-responsive clones and both of which showed a complete loss of this endogenous subunit on all annotated genes (**Figure 1J**). Finally, sequencing of nascent RNAs isolated from elongating polymerases complexes (Werner and Ruthenburg, 2015; Wuarin and Schibler, 1994; **Figure S1D**) further revealed a clear reduction of transcription (**Figure 1K**). This approach is well suited to evaluate transcription by all three types of polymerase (**Figure S1E**) and efficiently separates nascent chains from mature tRNAs (**Figure S1F**). Hence, these results confirm that the degradation protocol is efficient and sufficiently rapid to prevent notable changes in steady-state RNA levels (except for highly unstable species) and should therefore allow for the evaluation of primary effects of RPB1 loss on Pol III transcription. To our surprise the levels of RNAs in whole cell RNAs or in the polymerase-bound nascent RNA fraction were not dramatically affected by the DA-mediated depletion of Pol II. Pol II transcription is responsible for ∼60% of total RNA synthesis in mammalian cells (Pombo et al., 1999; Wansink et al., 1993). Hence if Pol II activity was indeed fully lost, one would expect a substantial reduction (at least 50%) in the levels of nascent RNAs per cells. Yet our measurements revealed a mere 23% reduction (**Figure 1L**), indicating a significant residual level of Pol II transcription in DA-treated cells.

**Figure 1.**
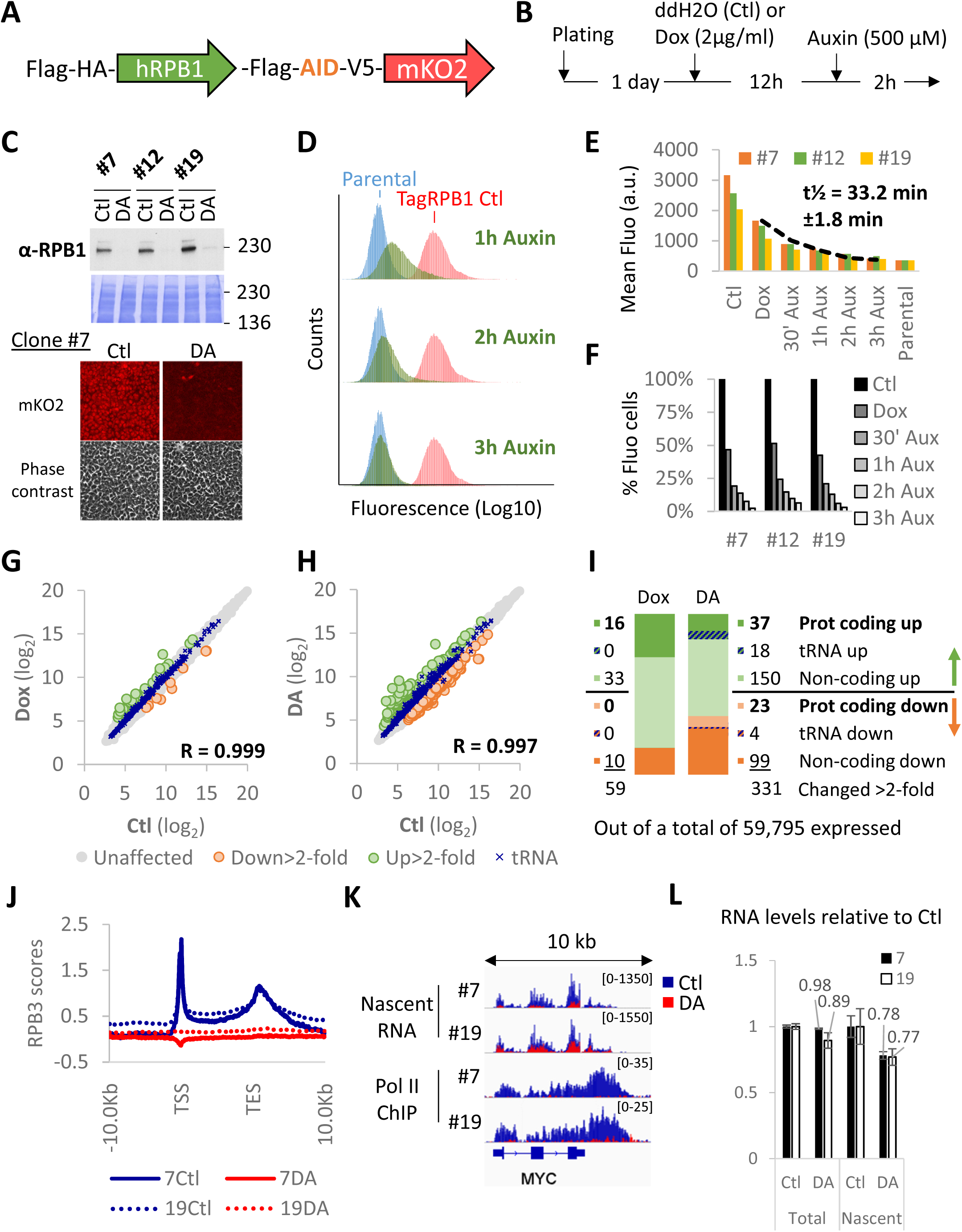
Establishment of a cell line for rapid and inducible depletion of RPB1. **(A)** Schematic representation of the tagged RPB1 construct expressing a full length human RPB1 cDNA flanked with an N-terminal Flag-HA tag and a C-terminal Flag tag followed by a minimal auxin-sensitive degron, a V5 epitope and an mKO2 orange fluorescent protein**. (B)** Standard protocol for RPB1 depletion. **(C)** RPB1 immunoblot of RIPA nuclear extracts prepared from all three clones used in this study and a representative example of a fluorescence microscopy analysis showing a DOX + auxin (DA)-dependent depletion of RPB1. **(D)** Time course flow cytometry experiment for clone #7 showing the progressive loss of fluorescence of DA-treated cells (in green). In red, untreated fluorescent cells. In blue, parental non-fluorescent cells. **(E)** Kinetics of fluorescence decay in all three clones. (**F)** % of remaining fluorescent cells at different time points during DA treatment for all 3 clones. **(G)** Correlation between gene expression levels in untreated (Ctl) and DOX-treated (DOX) pools of RNA prepared from all three clones. **(H)** Same as **(G)** but showing correlation between Ctl versus DA expression levels. **(I)** Number and type of genes affected >2-fold in the microarray analysis presented in **(G)** and **(H). (J)** Profile of RPB3 enrichment scores at all RefSeq genes for RPB1-expressing (Ctl) or RPB1-depleted (DA) clones #7 and 19#. TSS, transcription start site. TES, transcription end site. **(K)** Overlays of normalized nascent RNA and input-subtracted RPB3 ChIP-seq coverage at the *MYC* locus in RPB1-expressing (Ctl, blue) and RPB1-depleted (DA, red) cells for clones #7 and #19. **(L)** Quantification of RNA content in whole cell extract (Total) or nascent RNA fractions (Nascent) for clone #7 and #19 relative to Ctl.

### Dox + auxin-treatment converts a fraction of initiated Pol II into termination-incompetent CTD-less complexes

In an attempt to understand these discrepancies, we evaluated nascent RNA expression for the upregulated genes in the microarray (**Figure 2A**). To our surprise, most of these genes were expressed only at low levels prior to RPB1 depletion (see also **Figure 1H**). Importantly, however, virtually all of them were located downstream of highly expressed genes and the increase in their expression appeared to originate from termination site read-through from neighboring genes (**Figure 2A and 2C**). Importantly, termination defects were not observed for most genes (**Figure 1K and 2B**). MST1L lncRNA, the most upregulated gene (21-fold in the microarray analysis), is located downstream of a Pol II-transcribed U1 snRNA gene that presented a marked transcription termination defect upon DA treatment (**Figure 2C**), a feature shared by all Pol II-transcribed snRNA genes (**Figure 2D**). In addition, termination site read-through was also particularly pronounced at genes, such as UBC (Figure 2E and 2F), previously reported to display stable Pol II pausing (Chen et al., 2015). These results prompted us to reevaluate whether the tagged RPB1 construct was indeed fully degraded.

**Figure 2.**
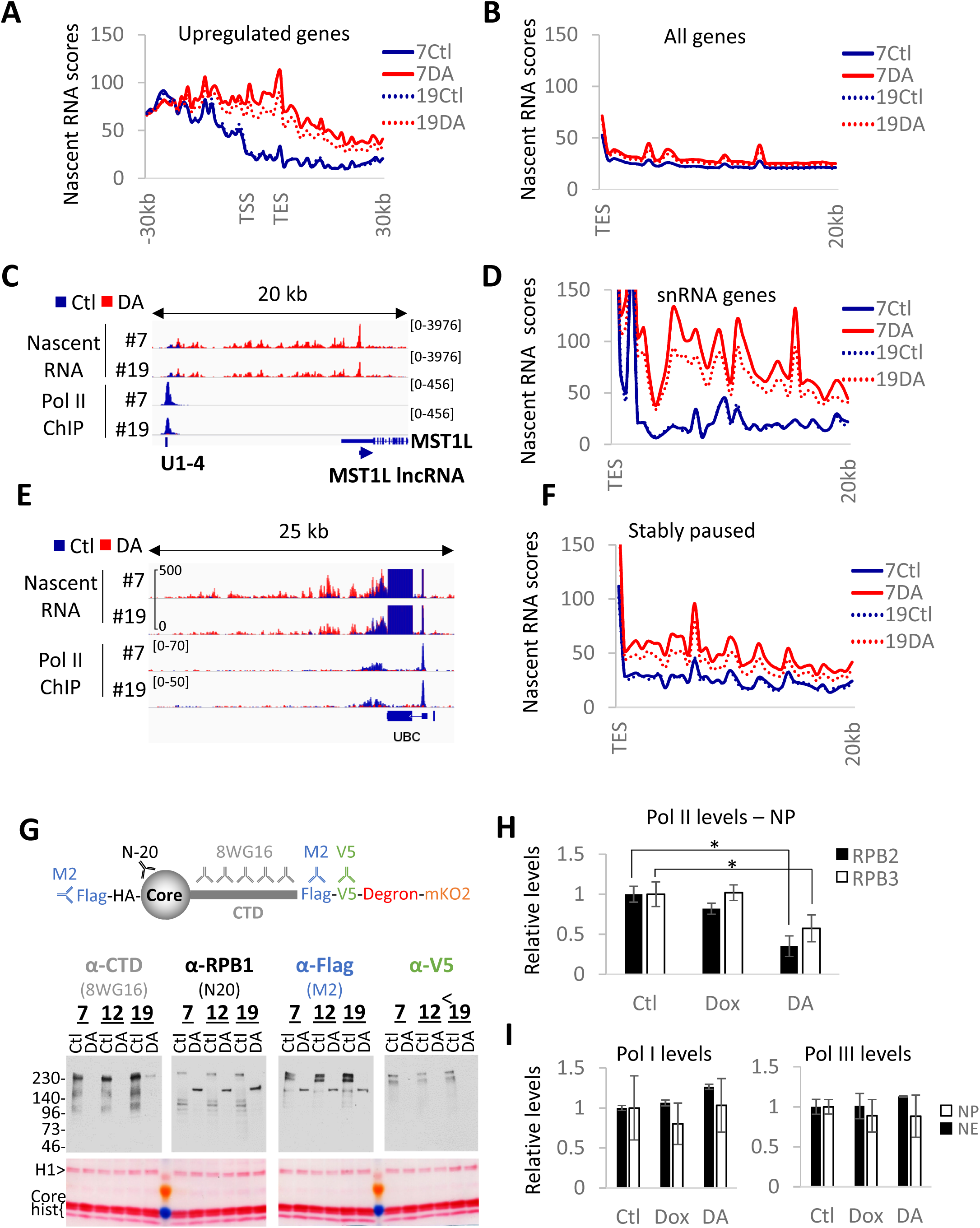
DA-treatment converts a fraction of initiated Pol II into termination-incompetent Pol IIB. **(A)** Profile of nascent RNA scores at Pol II genes upregulated in the microarray analysis presented in Figure 1H-I in RPB1-expressing (Ctl) or RPB1-depleted (DA) cells for clones #7 and 19#. TSS, transcription start site. TES, transcription end site. **(B)** Same as **(A)** but genome-wide profile (all RefSeq genes) of nascent RNA scores downstream of TES. **(C)** Overlays of normalized nascent RNA and input-subtracted RPB3 coverage at the MST1L lncRNA locus (anti-sense of MST1L), showing termination site read-through from the upstream snRNA U1-4 gene in DA treated clone #7 and #19. **(D)** Same as **(B)** but profile of nascent RNA scores downstream of all snRNA genes. **(E)** same as **(C)** at and downstream of the stably paused *UBC* gene showing termination site read-through for DA-treated clones #7 and #19 cells. **(F)** same as **(B)** but profiles of nascent RNA scores downstream of stably paused genes. **(G)** Schematic representation of tagged RPB1 protein and the different antibodies used to evaluate RPB1 degradation efficiency and immunoblot and corresponding Ponceau-stained membrane of chromatin pellets prepared from DA-treated and control cells. **(H)** Quantification of the levels of Pol II subunit expression in nuclear extracts and chromatin pellets **(see Figure S2B)**. **(I)** Same as **(H)** but quantification of Pol I and III expression levels. NP and NE correspond to, respectively, nuclear pellets and nuclear extracts. Data represent the average +/- SD. * indicates statistically significant difference (T-test; p<0.05).

Taking advantage of the multiple tags and different antibodies available for RPB1 (**Figure 2G**), we probed the chromatin-bound fraction (containing only initiated polymerases) isolated from untreated and DA-treated cells for RPB1 expression. Immunoblot analyses revealed the formation of truncated RPB1 lacking the CTD entirely (**Figure 2G, S2A and S2B**). Furthermore, immunoblotting for the endogenous Pol II subunits RPB2 and RPB3 confirmed that about a third of the transcribing Pol II was still present on the chromatin after RPB1 depletion (**Figure 2H and S2B**). In contrast, Pol I and Pol III levels remained unaffected (**Figure 2I and S2B**). Since Pol II with a CTD containing only 5 heptad repeats (out of a total of 52) is deficient for initiation in the context of chromatin (Lux et al., 2005; Meininghaus et al., 2000), the absence of truncated RPB1 in nuclear extracts (**Figure S2B**) suggests that the truncation occurred after initiation. Moreover, CTD-less Pol II (Pol IIB) is also deficient in termination (McCracken et al., 1997), and thus will remain associated with the chromatin. To confirm the absence of transcription initiation following DA treatment, serum-starved cells were stimulated with 20% fetal calf serum (FCS) following RPB1 depletion (**Figure S2C**). This treatment completely abolished induction of all serum-responsive genes (**Figure S2D and S2E**). Hence, the lack of RPB3 accumulation along target genes and the upregulation of non-expressed genes following RPB1 depletion are both explained by the conversion of a fraction of initiated Pol II into a form (CTD-free Pol IIB) that is competent for elongation but not for termination. However, the truncation was restricted to snRNA genes or genes displaying stable pausing.

### Pol II transcription interferes with RNA Pol III activity

Next, we postulated that a coupled auxin-mediated RPB1 depletion-nascent RNA expression analysis could provide insights into potential functions of Pol II at tRNA loci. Indeed, while complete depletion of Pol II would directly reveal its requirement at tRNA loci, this approach does not discriminate between the functions of transcribing, paused or promoter-bound Pol II molecules in Pol III-dependent transcription. However, we reasoned that the termination defect of CTD-less Pol II, which can easily be identified in nascent RNA-seq data, offered us a unique opportunity to examine directly the functional consequences of transcribing Pol II complexes at tRNA located downstream of a Pol II gene presenting termination site read-though. Finally, to identify all Pol II-sensitive tRNAs genes, we reanalyzed published nascent RNA-seq data (Werner and Ruthenburg, 2015) with wild-type HEK293 cells treated for 1 hr with the CDK9 inhibitor DRB, which blocks Pol II pause-release and therefore provides a reference where no Pol II transcription can occur at tRNA loci (**Figure 3A**). This analysis revealed that out of a total of 371 expressed tRNA loci, 55 were affected by DRB (44 upregulated and 11 downregulated at least 1.5-fold with an FDR <0.05). In our DA-treated HEK293 cells, only 8 of the 44 DRB-upregulated tRNA genes were also significantly increased by DA-mediated RPB1 depletion (**Figure 3B, S3A** and **Table S2**). As expected, tRNA loci that appeared upregulated because of high levels of Pol II read-through under DA conditions (**Figure S3A**) were not affected in DRB-treated cells. In addition, loci downregulated under DA conditions because they were located in transcribed Pol II genes were also not considered for further analysis (**Figure S3B**). We also confirmed that all DRB-sensitive tRNA genes were *bona fide* Pol III target genes in our cell lines by a ChIP-seq analysis of the RPC62 subunit of Pol III (**Figure S3C**).

**Figure 3.**
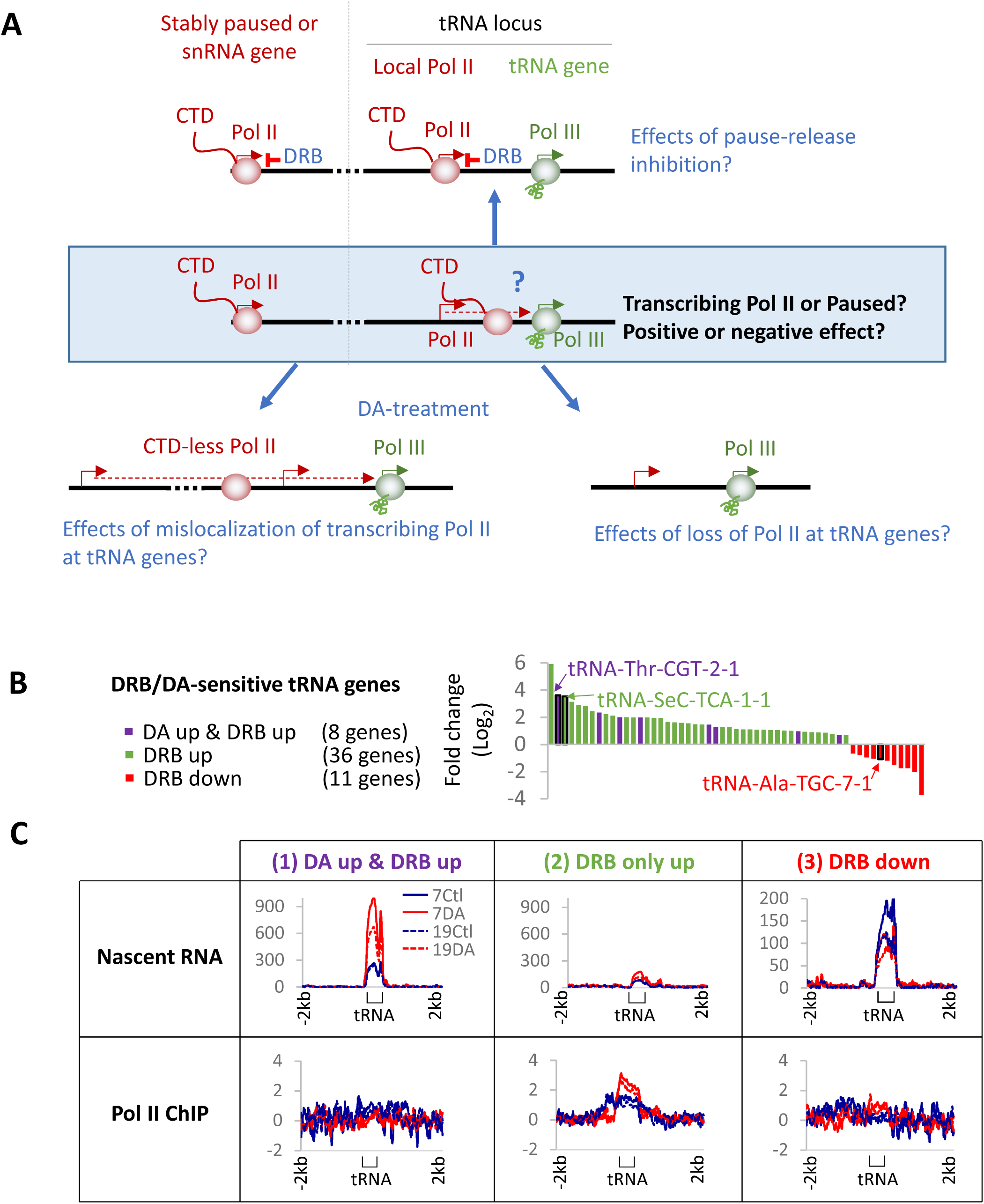
Pol II transcription interferes with RNA Pol III activity. **(A)** Schematic representation of the experimental approach and expected effects on tRNA gene transcription. See main text for details. **(B)** tRNA genes affected >1.5-fold at the nascent RNA level with a FDR<0.05 in DRB-treated HEK293 (data from (Werner and Ruthenburg, 2015)) ranked by Log_2_ fold change). tRNA genes affected significantly in DA-treated clones #7 and #19 cells are indicated by purple bars. Black rectangles identify the 3 model tRNA genes used in following experiments. **(C)** Profiles of nascent RNA and RPB3 enrichment scores at tRNA genes upregulated in both DA-treated and DRB-treated cells (1), upregulated in DRB only (2) or downregulated in DRB only (3) as defined in **(A).**

Given these observations, we then investigated Pol II occupancy at the three categories of tRNA loci defined in **Figure 3C**, namely: (1) those upregulated both in DA-treated cells and in DRB-treated cells at the nascent RNA level, (2) those upregulated only in DRB-treated cells and, finally, (3) those downregulated following DRB treatment (categories (1), (2) and (3) in **Figure 3C**). We did not observe Pol II densities at most tRNA loci (all categories) under control conditions (**Figure 3C,** 7Ctl and 19Ctl profiles). This suggests that Pol II does not accumulate at high levels near affected tRNA genes in HEK293. However, Pol II densities were observed at tRNA loci in the second category (upregulated in DRB only at the nascent RNA level) in DA-treated cells **(Figure 3C, (2))**. In addition, consistent with the presence of Pol II, clear evidence of active transcription upstream of most of these loci could also be observed upon DA treatment (**Figure 4A**, tRNA-SeC-TCA-1-1; **Figure S3D**, tRNA-Ser-GCT-5-1). Hence, the DRB-mediated increase in nascent tRNA levels is likely not caused by an indirect effect of Pol II transcription loss. Indeed, only a subset of tRNA genes was stimulated by DRB or DA treatment. Moreover, many DRB-induced tRNA genes remained unaffected in RPB1-depleted cells when Pol II transcription was maintained by CTD-less complexes (category (2)). Hence, the difference between category (1) genes that are upregulated in both DA- and DRB-treated cells and category (2) genes, upregulated only in DRB, lies in the maintenance of active Pol II transcription at these loci. In relation to the genes downregulated by DRB, none were affected to a statistically significant degree, relative to controls, by DA treatment (**Figure 4B, (3)**). These genes were nonetheless negatively impacted by RPB1 depletion but presented significant clonal variability even in non-treated cells.

**Figure 4.**
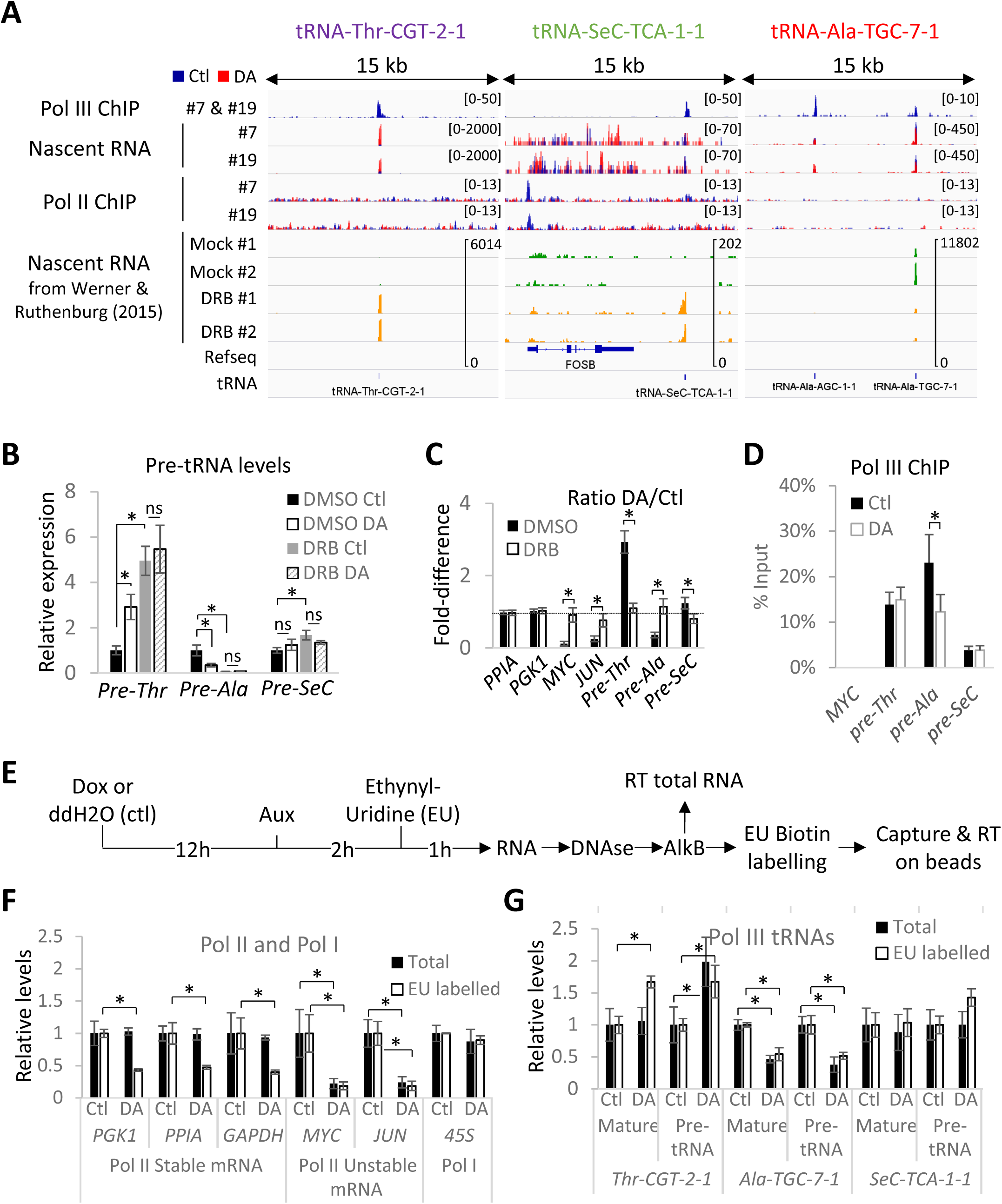
RPB1 depletion affects tRNA transcription rates. **(A)** Input-subtracted RPC62 ChIP-seq (overlay of clone #7 and #19 ChIP data), overlays of normalized nascent RNA and input-subtracted RPB3 ChIP-seq coverage in RPB1-expressing (Ctl) or RPB1-depleted (DA) clone #7 and #19 cells and nascent RNA coverage tracks of DRB-treated (green) and mock-treated (yellow) cells from Werner and Ruthenberg (2015). **(B)** Relative RNA expression levels in all three clones normalized to DMSO Ctl in RPB1-expressing (Ctl) or RPB1-depleted (DA) cells pre-treated with DRB or DMSO**. (C)** DA/Ctl ratios for data presented in **(B). (D)** RPC62 ChIP-qPCR analysis of chromatin preparations from clone #7 and #19 cells treated with auxin or DA. Data are represented as % of input. **(E)** Protocol used for the analysis of demethylated total and ethynyl-uridine (EU) labelled RNAs by RT-qPCR. **(F)** Relative expression levels of selected Pol I and Pol II RNAs in RPB1-expressing (Ctl) or RPB1-depleted (DA) cells for all 3 clones normalized to their levels under Ctl conditions. Data show the levels in total (Total) or metabolically labelled (EU-labelled) RNAs. **(G)** Same as **(F)** but for the three selected mature or precursor tRNAs. Data in **(B)-(G)** represent the average +/- SD. * indicates statistically significant differences, ns non-significant (T-test; p<0.05).

To further characterize the effects of Pol II inhibition/depletion on tRNA gene transcription in all three clones, we focused our attention on three model loci that are representative of the three categories of genes discussed above (**Figure 4C**): (1) tRNA-Thr-CGT-2-1 (pre-Thr), a DA-upregulated gene that is the second-most robustly overexpressed tRNA under DRB conditions; (2) tRNA-SeC-TCA-1-1 (pre-SeC), a gene that is regulated by type III external promoter and represents the third-most highly induced tRNA gene by DRB, but is unaffected by DA treatment. The lack of effect of the DA treatment at this locus correlates with the presence of transcribing, presumably termination incompetent, Pol IIB originating from the stably paused FOSB gene; (3) tRNA-Ala-TGC-7-1 (pre-Ala), a tRNA gene downregulated by DRB and one of the four tRNA genes downregulated in our microarray analysis of DA-treated cells (**Figure 1G and 1H**).

First, we confirmed that DA and DRB treatments mediated their effects via the same pathway by incubating cells for 1 hr with DRB prior to RPB1 depletion. DRB treatment, like RPB1 depletion, efficiently reduced the mRNA levels of unstable (*MYC, JUN*) Pol II transcripts and there was no additive effect of RPB1 depletion and DRB (**Figure 4C**). Similarly, DRB and DA treatments each showed robust increases in pre-Thr tRNA levels and decreases in pre-Ala tRNA levels with combined treatments being essentially equivalent to the somewhat stronger DRB effects (**Figure 4B and 4C**, DA/Ctl ratio = 1). Interestingly, in total RNA, pre-SeC tRNA levels remained unaffected following DA treatment but showed a statistically significant 1.7-fold increase with DRB. This latter observation confirmed that this tRNA is upregulated following DRB treatment but not by RPB1 deletion. Similar results could also be reproduced with the CDK9 inhibitor LDC00067 (LDC) and the initiation inhibitor Triptolide (Trip) pretreatment (**Figure S4A, S4B-and S4E).** These treatments confirmed that the effects observed are caused by the loss of Pol II transcription itself since degrading or inhibiting pause-release or initiation of Pol II have all the same effects. Moreover, since the triptolide treatment that was used induces the complete loss of Pol II at the time of gene expression analysis (**Figure S4A**), it appears very clear that all the expression changes in DA-treated cells at these loci are directly caused by modulation of Pol III activity. Finally, we evaluated the effects of RPB1 depletion on Pol III occupancy at these three loci by RPC62 ChIP-qPCR (**Figure 4D**). Pol III levels at the pre-Ala tRNA locus were reduced, confirming repression of this locus upon Pol II loss. In contrast, neither pre-Thr tRNA nor pre-SeC tRNA loci showed alterations in their Pol III levels. Since the density of a polymerase at a specific locus is determined by the ratio of the initiation frequency over the elongation rate (Ehrensberger et al., 2013), a higher tRNA synthesis rate without a change in the level of Pol III implies concomitant increases in both initiation and elongation rates (see Discussion).

Taken together, these results indicate that Pol II activity rapidly modulates Pol III transcription both positively and negatively at specific subsets of tRNA genes. Moreover, it is likely that Pol II transcription *per se*, rather than pausing or recruitment mediates these effects. Indeed, the exact same outcomes were obtained with drugs (LDC and DRB) that stall polymerases at pause sites and with a drug (Trip) that blocks initiation and induces complete loss of Pol II in these cells. The absence of detectable Pol II densities near regulated tRNA loci is incompatible with the presence of stably paused Pol II but not with an infrequent passage of transcribing Pol II complexes through these genes and an associated interference with the Pol III transcription cycle. A clear accumulation of Pol II near tRNA genes could only be observed when transcribing CTD-less complexes encountered tRNA genes strongly bound by Pol III (**Figure S3D**) – confirming that Pol III or its associated factors can slow down transcribing Pol II at tRNAs but not fully stop them as revealed by overlapping nascent RNAs (see also **Figure S6A**).

### RPB1 depletion affects tRNA transcription rates directly and indirectly

The fact that pre-Thr tRNA transcription can be enhanced without changes of Pol III occupancy or evidence of Pol II transcription prompted us to further evaluate whether these effects were indeed caused by modulation of Pol III activity or by changes in the stability or retention time of RNA precursors on chromatin. Thus, we monitored *de novo* synthesis of both stable tRNAs and their unstable precursors using metabolic labelling. Control or RPB1-depleted cells were incubated for 1 h with ethynyl-uridine (EU) (**Figure 4E**) and total RNA isolated from these cells was treated with an RNA demethylase (AlkB) to remove methyl groups that are known to interfere with RNA reverse transcription (Cozen et al., 2015; Zheng et al., 2015). Finally, EU-labelled RNAs were biotinylated, captured on streptavidin magnetic beads and analyzed by RT-qPCR. As expected, the synthesis of both stable and unstable mRNAs was substantially reduced (**Figure 4F**), but not abolished, by RPB1 degradation, consistent with the formation of elongating CTD-less complexes. The rapid loss of RPB1 did not alter EU incorporation either in Pol I transcripts or in most of the tested Pol III transcripts (**Figure 4F and S4F**). As expected, however, pre-Thr tRNA levels were increased in both total and EU-labelled RNA following DA treatment, consistent with an increase in transcriptional rates (**Figure 4G**). Furthermore, a significant Increase of EU-labelled mature tRNA could also be measured for tRNA-Thr (labelled pre-Thr tRNA, which is also detected by the primers used for the mature form, contributes only a small fraction (1/28) of the level measured; **Figures S4G and S4H**). In contrast, levels of both the pre- and mature Ala-tRNA were decreased in total RNA and EU labelled fractions following DA treatment, suggesting that this tRNA is unstable in HEK293 cells (**Figure 4G**). Finally, increased levels of EU incorporation could be observed for pre-SeC tRNA but not for its mature form, consistent with the changes detected at the nascent RNA level. Hence, these changes mainly reflect higher EU incorporation into readthrough transcripts that are synthesized by CTD-less complexes and cannot be processed and remain associated with the chromatin.

To further confirm that these effects reflected changes in transcriptional rates rather than stability/processing of the tRNA precursors, control and DA-treated cells were incubated with ActD to fully block transcription by all RNA polymerases and pre-tRNA levels were measured at different time points. The decay rates of all three tested pre-tRNAs were not significantly affected by RPB1 depletion (**Figure S4I and S4J**), confirming that changes in expression levels mirrored alterations in Pol III transcriptional rates. In addition, changes in pre-tRNA synthesis rates following RPB1 depletion were also independent of protein synthesis, as revealed by pre-incubation with the translation inhibitor cycloheximide (CHX; **Figure S4K**). Hence, stimulation of pre-Thr tRNA transcription following RPB1-depletion is likely direct. However, in the case of the downregulated pre-tRNAs, these experiments did not formally rule out an indirect effect such as loss of an unstable activator(s). Except for *MYC*, none of the genes encoding known Pol III factors or activators were among the 23 protein-coding genes affected in DA-treated cells (**Figure 5A and Table S3**). Indeed, MYC has been shown to exert a positive stimulatory role on Pol III transcription via recruitment of GCN5 and TRRAP (Kenneth et al., 2007). Since MYC levels rapidly decreased during the DA-treatment (**Figure 5A and 5B**), we decided to evaluate the MYC contribution to pre-tRNA expression by transfecting cells with a siRNA pool targeting *MYC* (siMyc) for 36 hrs prior to RPB1 depletion (**Figure S5A**). While *MYC* knock-down at the mRNA level was very efficient (**Figure S5C**), it also interfered with Dox-mediated induction of the TIR1 ubiquitin ligase and, as a consequence, impaired RPB1 degradation in a large number of cells (**Figure S5B and S5C**). As a consequence, pre-Thr tRNA appeared less induced by the DA-treatment but its expression in RPB1-expressing cells remained unaffected by MYC knock-down (**Figure 5C**). Similarly, Pre-SeC tRNA levels were not significantly altered by siRNA-mediated depletion of MYC. In contrast, pre-Ala tRNA levels were reduced by *MYC* siRNAs to the same extent that they were reduced in DA treated cells. Although the decrease in pre-Ala tRNA levels did not strictly parallel the extent of MYC mRNA reduction, this tRNA gene nonetheless appears to be sensitive to MYC levels. Hence, the downregulation of pre-Ala tRNA observed after RPB1 depletion seems to be caused, at least in part, by the loss of MYC and thus more likely reflects indirect effects. Interestingly, the three clones displayed different levels of expression of MYC protein (**Figure 5A and 5B**), which correlates with the difference in the levels of nascent RNAs observed in control versus treated cell for the tRNA loci downregulated by DRB (**Figure 3C (3)).**

**Figure 5.**
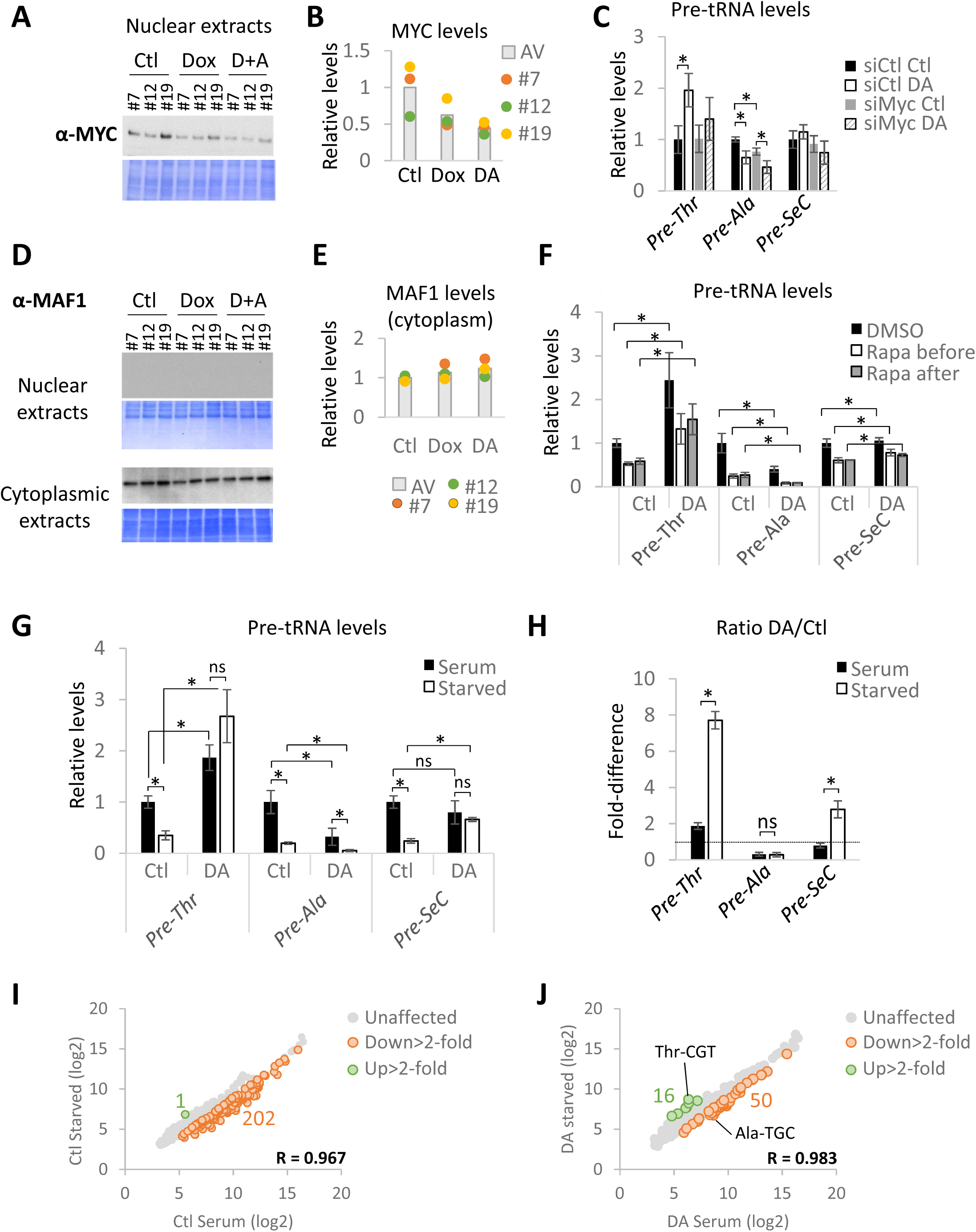
Pol II activity regulates tRNA transcription directly and indirectly and is essential for the repression of most tRNA genes in serum-deprived cells. **(A)** MYC immunoblot and corresponding Coomassie-stained membrane of nuclear extracts prepared from untreated (Ctl), Dox-treated and DA-treated cells for all three clones. **(B)** Quantification of data shown in **(A).** Histogram represents the average expression. **(C)** Relative pre-tRNA expression levels normalized to siCtl-treated cells for all 3 clones transfected with a non-targeting (siCtl) or *MYC*-targeting pool of siRNAs (siMyc) prior to RPB1 depletion. **(D)** MAF1 immunoblots and corresponding Coomassie-stained membrane of nuclear and cytoplasmic extracts. **(E)** MAF1 protein levels in cytoplasmic extracts prepared from RPB1-expressing or RPB1-depleted cells for all 3 clones. Histograms represent the average expression levels **(F)** Relative pre-tRNA levels normalized to Ctl DMSO levels in RPB1-expressing (Ctl) or RPB1-depleted (DA) cells for all three clones treated with Rapa prior to or after auxin addition**. (G)** Relative pre-tRNA levels normalized to Ctl cells kept in serum in RPB1-expressing (Ctl) or RPB1-depleted (DA) cells for all three clones cultured in the presence (black) or absence (white) of serum prior to and during RPB1-depletion. **(H)** DA/Ctl ratios for data shown in **(G). (I)** Microarray analysis of pools of RNA prepared from all three clones cultured in the presence (serum) or absence of serum (starved) for 20hrs. **(J)** Same as **(I)** but for cells treated with DA. Data in **(C), (F), (G) and (H)** represent the average +/- SD. * indicates statistically significant differences, ns non-significant (T-test; p<0.05).

### Pol II activity is essential for the repression of tRNA genes in serum-deprived cells

The fact that Pol II transcription loss did not affect the transcription of all tRNAs suggests a direct participation of Pol II in the modulation of Pol III activity at specific loci rather than an alteration of the levels or activity of a general regulator. However, as shown above, changes in MYC levels specifically modulated pre-Ala expression. By analogy, mammalian and yeast tRNAs display different sensitivities to the global Pol III repressor MAF1 (Cieśla et al., 2007; Orioli et al., 2016; Turowski et al., 2016). In human cells, MAF1 is essential for Pol III repression in response to unfavorable growth conditions, such as serum starvation (Michels et al., 2010). This Pol III repressor was also shown to exert a negative control under basal conditions, at least in IMR90 and HEK293 cells (Reina et al., 2006). Hence, we assessed the effects of RPB1 degradation on pre-tRNA expression in MAF1-depleted cells. First, we confirmed that MAF1 protein levels and subcellular localization were not affected by the DA treatment (**Figure 5D and 5E**). To investigate whether repression by Pol II transcription is achieved via MAF1 or whether both exert independent negative effects, we manipulated MAF1 activity instead of its levels using rapamycin (Rapa). This mTORC1 inhibitor was shown to prevent MAF1 phosphorylation and inactivation in mammalian cells (Michels et al., 2010; Shor et al., 2010). Cells thus were treated either with DMSO or with rapamycin (Rapa) prior to or after RPB1 depletion (**Figure S5D**). While Rapa could repress all three pre-tRNAs (**Figure 5F**), it had no impact on the stimulation of pre-Thr tRNA transcription or the repression of pre-Ala tRNA transcription by RPB1 depletion and did not change the response of pre-SeC tRNA transcription to RPB1 degradation. These effects were also not affected by inverting the order of RPB1 depletion and Rapa addition (**Figure S5E**). These results indicate that MAF1 and Pol II act via independent mechanisms. Interestingly, the relative levels of pre-Thr tRNAs in Rapa plus DA-treated cells was similar to the level of pre-Thr tRNA in cells cultured in control conditions (Ctl, DMSO) (**Figure 6F**). Hence, loss of Pol II transcription cancelled the repression imposed by Rapa.

**Figure 6.**
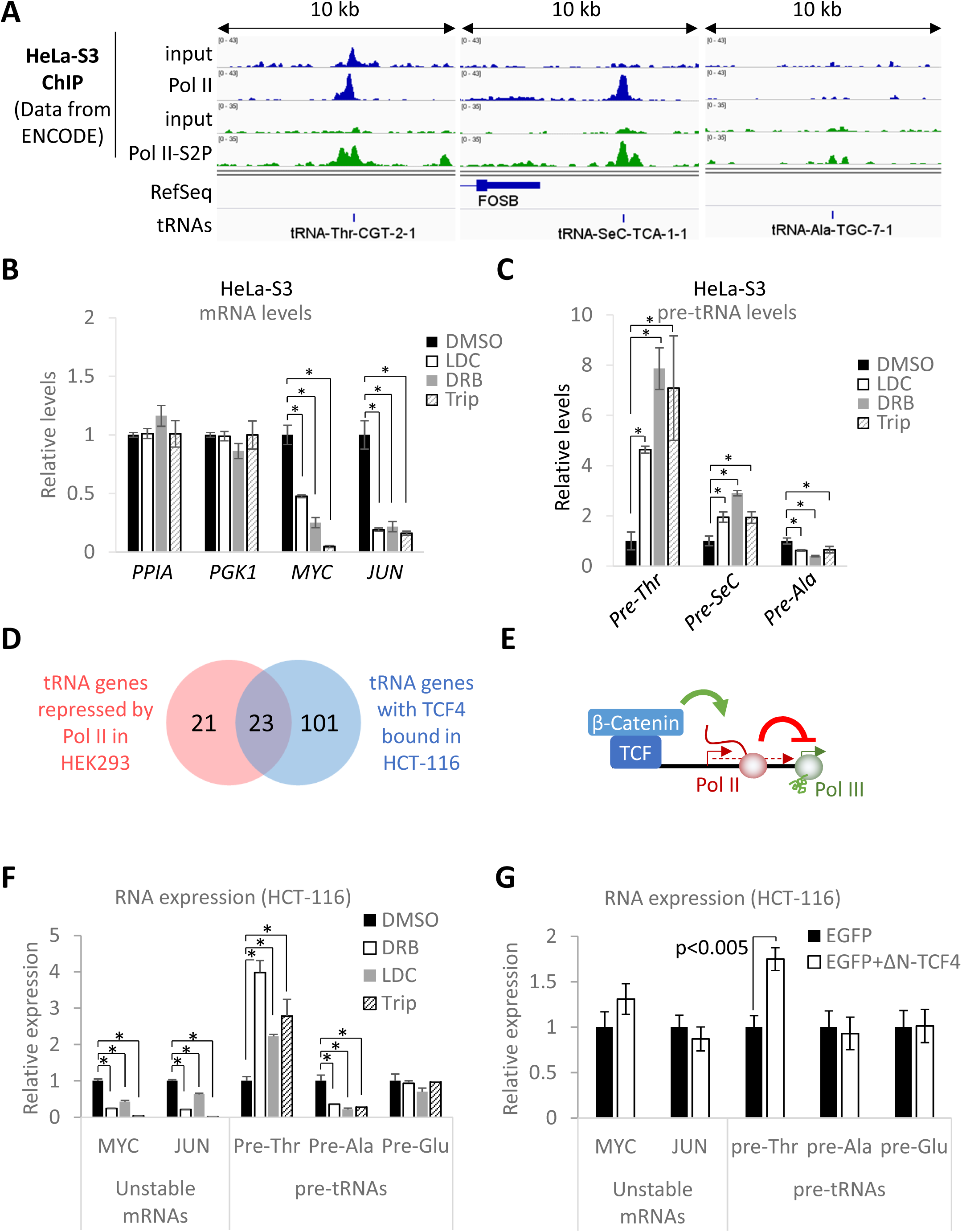
Pol II repression of tRNA genes is not limited to HEK293 cells and likely occurs directly from local Pol II promoters. **(A)** ChIP-seq coverage for total and Ser2-phosphorylated RPB1 and corresponding input (data from ENCODE) in HeLa S3 cells at three model pre-tRNA loci. **(B)** Relative mRNA levels in HeLa S3 cells treated for 1hr with LDC, DRB and Trip normalized to levels in DMSO-treated cells. **(C)** Same as **(B)** but showing normalized relative pre-tRNA levels. **(D)** Venn diagram showing that half of the tRNA genes up-regaluted by DRB in HEK293 cells are bound by TCF4 (within 500 bps from the tRNA genes) in HCT116 cells. **(E)** Model of β-catenin/TCF4-mediated repression of tRNA genes sensitive to Pol II repression. **(F)** same as **(B, C)** but for HCT-116 cells. **(G)** Relative mRNA and pre-tRNA levels in HCT-116 cells transiently cotransfected with plasmids expressing a dominant-negative TCF4 construct (ΔN-TCF4) and an EGFP or just an EGFP (control). Data in **(B), (C), (F) and (G)** represent the average +/- SD. * indicates statistically significant differences (T-test; p<0.05).

To evaluate the full extent of this effect of RPB1 depletion on Rapa-mediated activation of MAF1, which is normally activated under low growth conditions, we measured pre-tRNA expression in cells deprived of serum for 18 hrs prior to RPB1 depletion (**Figure S2C**). To our surprise, pre-Thr tRNA levels in serum-starved cells were up to 8-fold higher following RPB1 depletion, indicating that loss of Pol II activity for just 2 hrs was sufficient to fully reverse the normal low serum-mediated repression of this gene (**Figure 5G and 5H**). Similarly, RPB1 depletion in serum-starved cells restored pre-SeC tRNA expression to a level comparable to that observed in cells at high serum (**Figure 5G and 5H**). In contrast, the normal repression of pre-Ala tRNA transcription in serum-starved cells was enhanced by RPB1 depletion (**Figure 5G and 5H**, DA/Ctl <1). Strikingly, a microarray analysis performed with pools of RNAs prepared from all three clones treated or not with DA revealed that the transcriptional response of a large majority of tRNA genes affected by serum starvation in Ctl treated cells remain unaffected in serum starved DA-treated cells (202 vs 50; compare **Figure 5I to 5J**). Finally, serum stimulation of deprived cells induced a rapid and robust induction of all three pre-tRNAs in the presence and absence of Pol II (**Figure S5F**). However, the kinetics of these inductions were different for the pre-Thr and pre-SeC tRNAs, which both displayed faster responses to serum stimulation in the absence of Pol II – as revealed by the slopes of the inductions in the first 30 min (**Figure S5F**, red triangles). Altogether these results indicate that RNA Pol II transcription plays an essential role in the regulation of the transcriptional response of most tRNA genes in response to environmental cues.

### Pol II transcription can repress Pol III function directly from local promoters

Data accumulated so far are compatible with a model in which Pol II transcription can modulate Pol III activity directly via transcriptional interference or indirectly via loss of an unstable (co)activator(s). However, we were not able to show unambiguously that Pol II transcription occurs at tRNA genes that are upregulated by transcriptional inhibitors and by RPB1 depletion. Moreover, it also is not clear if these effects are restricted to HEK293 cells. Since Pol II was observed near 70% of all Pol III-bound tRNA loci in HeLa cells (Oler et al., 2010), we analyzed the presence of serine 2-phosphorylated (S2P) Pol II, a CTD modification associated with elongating complexes (Komarnitsky et al., 2000), at these loci in HeLa S3 cells using available ENCODE datasets. In this cell line, clear densities of Pol II and S2P-Pol II could be observed at both pre-Thr and pre-SeC tRNA loci but not at the pre-Ala tRNA locus (**Figure 6A**). If Pol II directly interferes with Pol III activity, one could expect that higher levels of Pol II should be correlated with greater repression. We tested this hypothesis using DRB, LDC and Trip treatments and measured pre-tRNA expression levels in HeLa S3 cells (**Figure 6B**). As expected, a 2-hr treatment with any of these inhibitors robustly stimulated expression of pre-Thr (up to 8-fold) and pre-SeC (up to 3-fold) tRNAs, but not pre-Ala tRNA (which was downregulated in all three cases as observed in HEK293 cells) (**Figure 6C**). These data support the hypothesis that tRNA repression by transcribing Pol II complexes is both direct and not restricted to HEK293 cells. Moreover, analysis of nascent RNA levels in DRB- and DA-treated HEK293 cells at a Pol III-transcribed *MIR* element located in the first intron of the Pol III subunit gene *POLR3E* (Yeganeh et al., 2017) showed that loss of Pol II transcription is accompanied by a clear upregulation of the Pol III gene (**Figure S6A**), further confirming that Pol II directly controls Pol III activity at specific loci via transcriptional interference. Next, we examined whether the tRNA-Thr locus displayed characteristics of a Pol II promoter by examining available ENCODE datasets for general transcription factor (GTFs) as well as gene specific regulators in all available cell lines (**Figure S6B**). This analysis revealed the presence of GTFs (such as TFIIF and TFIID) as well as numerous transcription factors in the immediate vicinity of this Pol III gene. Among them, the Wnt regulated factor TCF4 in HCT-116 cells was further found to localize in close proximity of half of all the DRB-sensitive tRNA loci identified (**Figure 6D**). Hence, if our model is correct, one can make the prediction that TCF4 should be involved in the repression of the tRNA-Thr locus by stimulating Pol II transcription (**Figure 6E**). To test this hypothesis, we first confirmed that the pre-Thr and pre-Ala tRNAs were indeed also regulated by Pol II in these cells using transcriptional inhibitors (**Figure 6F**). These results further confirmed the generality of Pol II repression in yet another cell line. Next, we assessed the function of TCF4 by transfecting a plasmid expressing a dominant-negative TCF4 (ΔN-TCF4) (Korinek et al., 1997). Despite a relatively low transfection efficiency (**Figure S6C),** cells expressing the dominant-negative constructs (**Figure S6D**), contributed sufficiently to the analyzed pool of RNAs to reveal a highly specific upregulation of tRNA-Thr without any effects on the tRNA-Ala loci and with no changes in MYC mRNA expression (**Figure 6G**). These results strongly indicate that Pol II, recruited specifically at local promoters by Pol II-specific transcriptional activators can regulate Pol III-dependent tRNA transcription via transcriptional interference.

## Discussion

The relative simplicity in the Pol III transcriptional machinery allows an efficient global regulation of RNA species involved in protein biosynthesis to match cellular demands. However, several lines of evidence suggest that individual tRNAs are specifically modulated in response to internal and external cues to support specific roles in a large number of biological processes (see (Rak et al., 2018). How gene-specific transcription of tRNAs genes is established remained largely unexplored in mammalian cells. During the past 10 years, numerous ChIP-seq studies revealed a surprising genomic colocalization of Pol II near Pol III genes (Barski et al., 2010; Canella et al., 2012; Moqtaderi et al., 2010; Oler et al., 2010; Raha et al., 2010; Yeganeh et al., 2017). However, the functional significance of these observations is difficult to address using conventional techniques since the mature forms of Pol III RNAs are very stable, highly abundant and difficult to measure.

To overcome these issues and explore functionally the connections between Pol II and Pol III transcription events, we designed cell lines in which the rapid depletion of RPB1 can be induced by the addition of auxin. Surprisingly, loss of Pol II resulted in rapid and severe consequences on the transcription of specific tRNA genes as revealed by nascent-RNA sequencing, pre-tRNA level measurement and metabolic labeling, which directly assess the Pol III transcriptional output. Our approach unambiguously identified Pol II as a direct gene-specific regulator of tRNA transcription by Pol III. In addition, fortuitously, these cell lines also allowed us to assess the effects of Pol II transcription at tRNA loci as a result of a partial truncation of the CTD in Pol II complexes at snRNA and stably paused genes.

### Identification of Pol II as a gene-specific regulator of tRNA transcription by Pol III

Rapid depletion of RPB1 by doxycycline and auxin (DA) treatment as well as pharmacological inhibition of Pol II revealed that Pol II transcription can modulate the synthesis of specific pre-tRNAs either positively or negatively (summarized in Figure 7C). Our results suggest that the decrease in expression of Pol III targets observed following RPB1 depletion (current results) or in α-amanitin treated cells (Barski et al., 2010; Listerman et al., 2007; Raha et al., 2010) is most likely indirect and results from the loss of unstable Pol III regulators such as MYC, previously shown to be a positive regulator of Pol III transcription (Kenneth et al., 2007). In contrast, repression of certain tRNA loci by Pol II is direct and does not require *de novo* protein synthesis. Loss of Pol II transcription was accompanied by higher levels of nascent tRNAs associated with repressed loci and by increased incorporation of EU into corresponding precursors and mature tRNAs. Moreover, the stability of the precursors tested was not affected by RPB1 depletion, confirming modulation of the synthesis of these pre-tRNAs. Further, the increase in tRNA transcription caused by Pol II inhibitors was not observed in RPB1-depleted cells when Pol II transcription was maintained by the termination-deficient, CTD-free Pol IIB formed at neighboring Pol II genes. To our surprise, we also found that Pol II transcription was essential to sustain repression of many tRNA genes during rapamycin treatment, which inhibits mTORC1 and thereby prevents MAF1 inactivation (Michels et al., 2010; Shor et al., 2010), and during serum starvation. Hence, MAF1 requires Pol II transcription to fully repress most tRNA genes. Altogether our results support a direct and essential role of Pol II in the repression of tRNA transcription not only in basal conditions but also in response to environmental cues and signaling pathways as diverse as the Wnt pathway.

**Figure 7.**
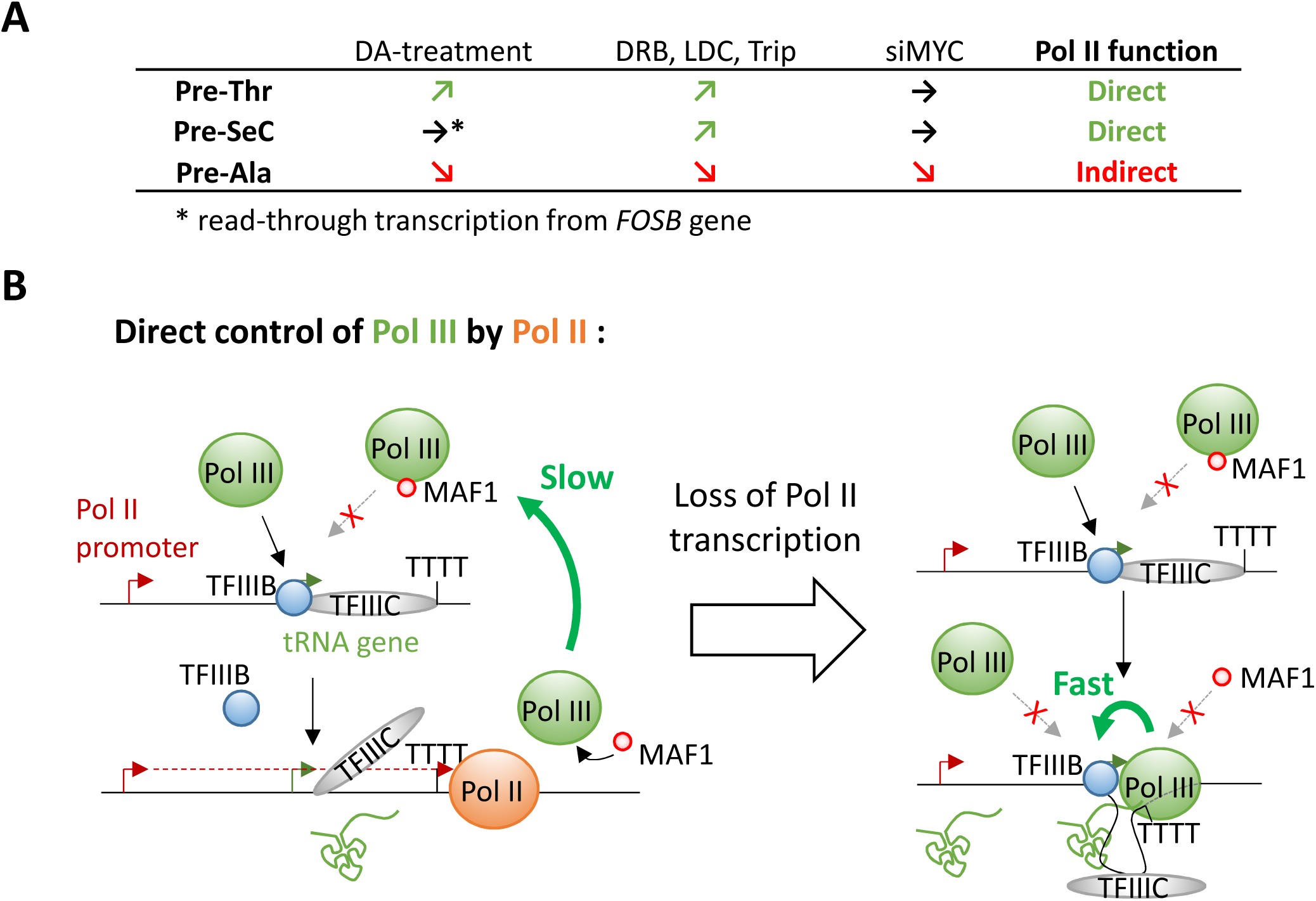
A model for direct repression of Pol III activity at tRNA genes by Pol II transcription. **(A)** Summary of the effects of different treatments on the expression of three pre-tRNAs**. (B)** Model for the Pol II regulation of tRNA transcription rates. See Discussion for details.

### A model for Pol III repression by transcriptional interference

Following the first round of transcription, terminating Pol III was found to be committed to reinitiation on the same template for multiple rounds without being released in solution in a process referred to as facilitated recycling (Dieci and Sentenac, 1996; Dieci et al., 2013). Polymerases engaged in such hyper-processive cycles transcribe their targets at much higher rates (between 5- to 10-fold) and are resistant to MAF1 (Cabart et al., 2008). Structural studies showed that MAF1 binds the same Pol III interface bound by TFIIIB, thereby preventing Pol III recruitment to cognate promoters (Vannini et al., 2010). Interestingly, reconstitution of Pol III elongating complexes in the presence of promoter-bound TFIIIB revealed that Pol III can interact with TFIIIB even in the gene body and likely during the entire transcription cycle (Han et al., 2018). This observation provides a mechanistic basis not only for facilitated recycling but also for the resistance of engaged polymerases to MAF1 in vitro. Although it still is not clear if facilitated recycling occurs *in vivo* (Arimbasseri et al., 2014), electron microscopic studies of 5S loci in yeast revealed an apparent interaction between the promoter and the terminator of some of these genes – an observation compatible with the existence of facilitated recycling in living cells (French et al., 2008). Hence, if recycling does occur *in vivo*, polymerases engaged in such hyper-processive loops would require an external player to break the cycling and allow MAF1 to establish repression. Based on our observations, transcribing Pol II passage through regulated tRNAs could assume this function by displacing, or at least interfering with, TFIIIB-Pol III interactions, thus allowing Pol III to interact with MAF1 and preventing subsequent recruitment to (re)assembled promoters (Figure 7D). This model further explains the apparent reversion of tRNA repression following Pol II loss during serum starvation. Indeed, since MAF1 is 4- to 10-fold less abundant than Pol III in mammalian cells (Orioli et al., 2016), it is very likely that every tRNA gene will encounter at least one MAF1-free Pol III complex during the 2-hr period of RPB1 depletion in our analyses, thereby allowing facilitated recycling to be reestablished. In addition, since the dwell time of reinitiating polymerases at promoters and terminators has to be shorter than the dwell time of newly initiating Pol III, the proposed model also explains how higher rates of transcription can be achieved without increased Pol III occupancy at the pre-Thr tRNA locus analyzed here – providing that elongation also occurs at a higher speed.

Hence, the presence of Pol II peaks close to tRNAs genes in some cell lines but not others could simply reflect cell line/tissue specific differences in nearby Pol II promoter activity. Higher local Pol II initiation rates would increase the chance of accumulation of elongating complexes at tRNA genes as they are slowed down by Pol III or its associated factors. Indeed, the strength of pre-Thr or pre-SeC tRNA stimulation following pharmacological inhibition of Pol II correlated with the higher levels of Pol II seen in HeLa cells. Our analysis also revealed that transcription factors such as the T Cell factor 4 (TCF4) can indirectly repress the pre-Thr-CGT-2-1 gene via Pol II in HCT-116 cells. Hence, systematic identification of these promoters and the specific regulators involved in their regulation will provide insights into the regulation of particular tRNAs in response to environmental cues, adverse conditions or pathologies. Such a transcriptional control is meaningful only if a sufficient number of genes encoding a particular tRNA isodecoder (anticodon) are co-regulated. Interestingly, more than 50% of all expressed Ala-ACG, Arg-TCG, iMet, SeC and Ser-CGA and -GCT tRNAs appeared to be enhanced upon DRB treatment (**Figure S7**), which therefore could potentially affect the abundance of proteins relying on these tRNAs for their translation. This is particularly interesting in the case of initiator methionine (iMet), which is essential for translation initiation and whose levels have been shown to control body size and developmental timing in Drosophila (Rideout et al., 2012). Moreover, an upregulation of this tRNA by only 1.4-fold was found to induce a global reprogramming of tRNA expression and to increase proliferation in human epithelial cells (Pavon-Eternod et al., 2013). However, the fact that most tRNA genes did not react to Pol II inhibition or depletion, or that some were even downregulated, suggests that the proposed mechanism is not general. It would therefore be of particular interest to evaluate whether these genes are capable of facilitated recycling *in vivo* and, if so, how these cycles are interrupted in adverse conditions. Alternatively, the dependency of pre-Ala tRNA levels on MYC levels may also indicate that high rates of transcription are perhaps achieved through stimulation of *de novo* initiation frequencies at this tRNA gene.

## Acknowledgements

We thank Sohail Malik, Takashi Onikubo, Evelina Tutucci and Ueli Schibler for their comments on the manuscript. This work was supported by NIH grants CA129325, DK071900 and CA202245 to R.G.R. A.G. was supported by a Swiss National Science Foundation Early Mobility Fellowship (P2GEP3_151952) and by a Human Frontier Science Program Long-Term Fellowship (LT001083/2014). K.I. was supported by a NCI T32 grant (CA009673) and a JSPS postdoctoral fellowship for research abroad.

## Author Contributions

A.G. designed and performed experiments, analyzed and interpreted data and wrote the manuscript. K.I. performed FACS experiments. C-S.C. performed ChIP experiments. R.G.R supervised the project and wrote the manuscript. All authors read the manuscript.

## Supplementary figure legends

**Figure S1.**
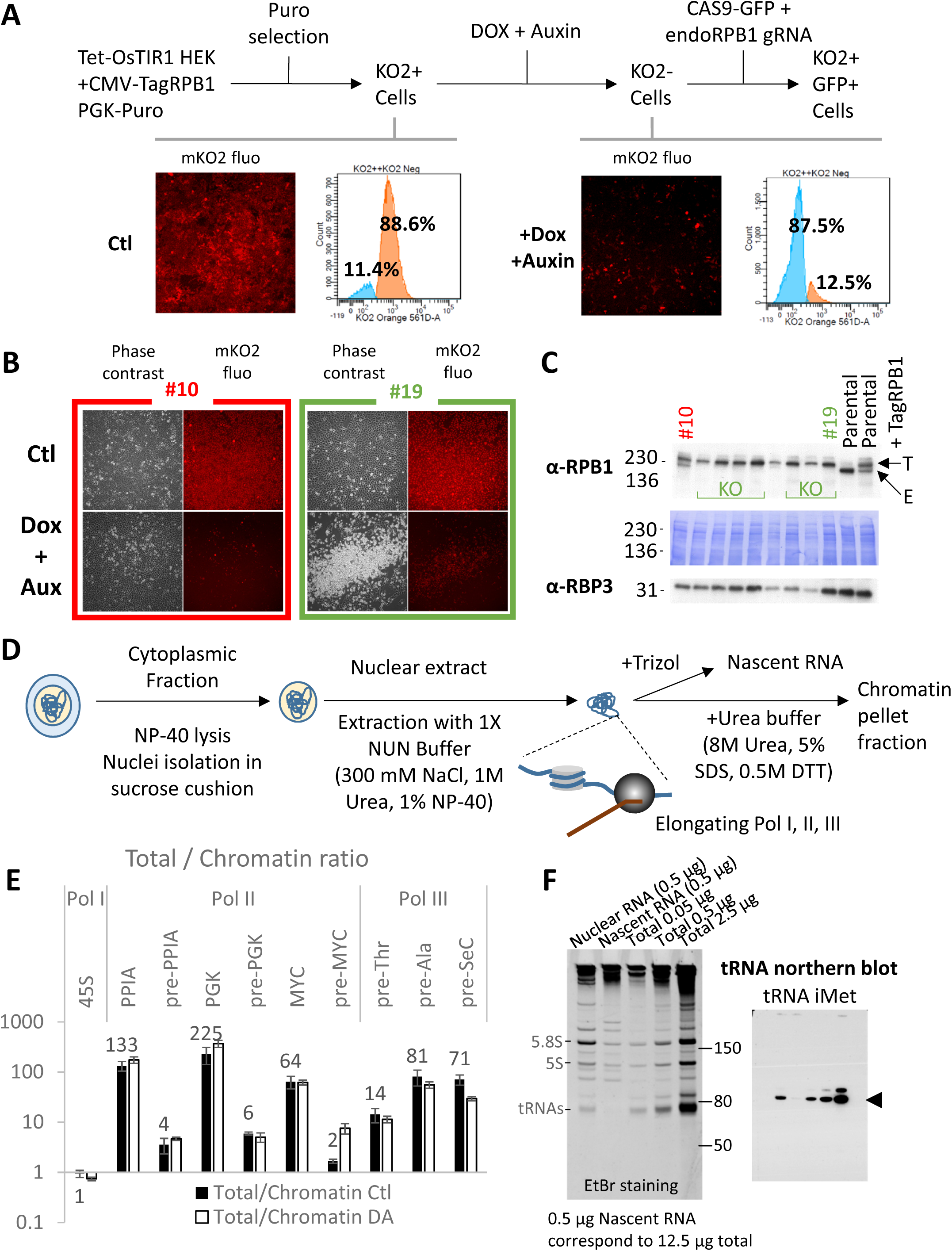
Related to Figure 1. **(A)** A construct encoding a full length RPB1 tagged with a fluorescent protein and auxin-degron was first stably transfected into an HEK293 cell line expressing a DOX-inducible OsTIR1 from the AAVS1 locus (Natsume et al., 2016). Dox and auxin (DA)-sensitive mKO2-expressing cells (non-fluorescent cells sorted in bulk after DA treatment) were then transfected with a plasmid expressing a guide RNA targeting selectively the endogenous RPB1 gene and a Cas9-GFP construct. Cells positive for both mKO2 and GFP were finally sorted as single clones. 146 clones were first screened for substantial cell death after prolonged treatment with DA, indicating inactivation of all endogenous copies of RPB1 as shown in **(B)** for a clone depending (#19, green) or not (clone #10, red) on the tagged RPB1 for survival. The data also show efficient depletion (loss of mKO2 fluorescence level) of the tagged construct in presence of auxin with massive cell death (#19) or no effect on cell viability (#10). 47 clones (32%) passed this first round of selection and the absence of endogenous RPB1 was confirmed by immunoblot using RIPA extracts prepared from isolated nuclei as shown in **(C)** for a few representative clones. E indicates the position of the endogenous RPB1 and T the position of the tagged RPB1. **(D)** Protocol used for the preparation of cytoplasmic, NUN-nuclear and pellets protein extracts as well as nascent RNA isolation. **(E)** Ratio of different RNA levels in total RNA versus chromatin bound fraction in RPB1-expressing (Ctl) or RPB1-depleted (DA) cells for clones #7, #12 and #19**. (F)** Left panel, EtBr staining of a denaturing polyacrylamide gel loaded with RNAs prepared from isolated nuclei (Nuclear RNA), NUN-extracted chromatin pellets (Nascent RNA) and increasing amount of total RNA (Total). Right panel, tRNA-iMet Northern blot showing complete depletion of mature tRNAs in the nascent RNA fraction. Note that 0.5 μg Nascent RNA corresponds to 12.5 μg of total RNA. This approach is well suited to evaluate transcription by all three types of polymerases. Indeed, while the 45S rRNA precursor is fully retained in the nascent RNA fraction, pre-mRNAs are clearly enriched compared to their mRNA counterpart as shown in **(E)**. Pre-tRNAs however are likely not processed co-transcriptionally and are expected to be released from the polymerase before excision of leader and trailer sequences (Arimbasseri, 2018).

**Figure S2.**
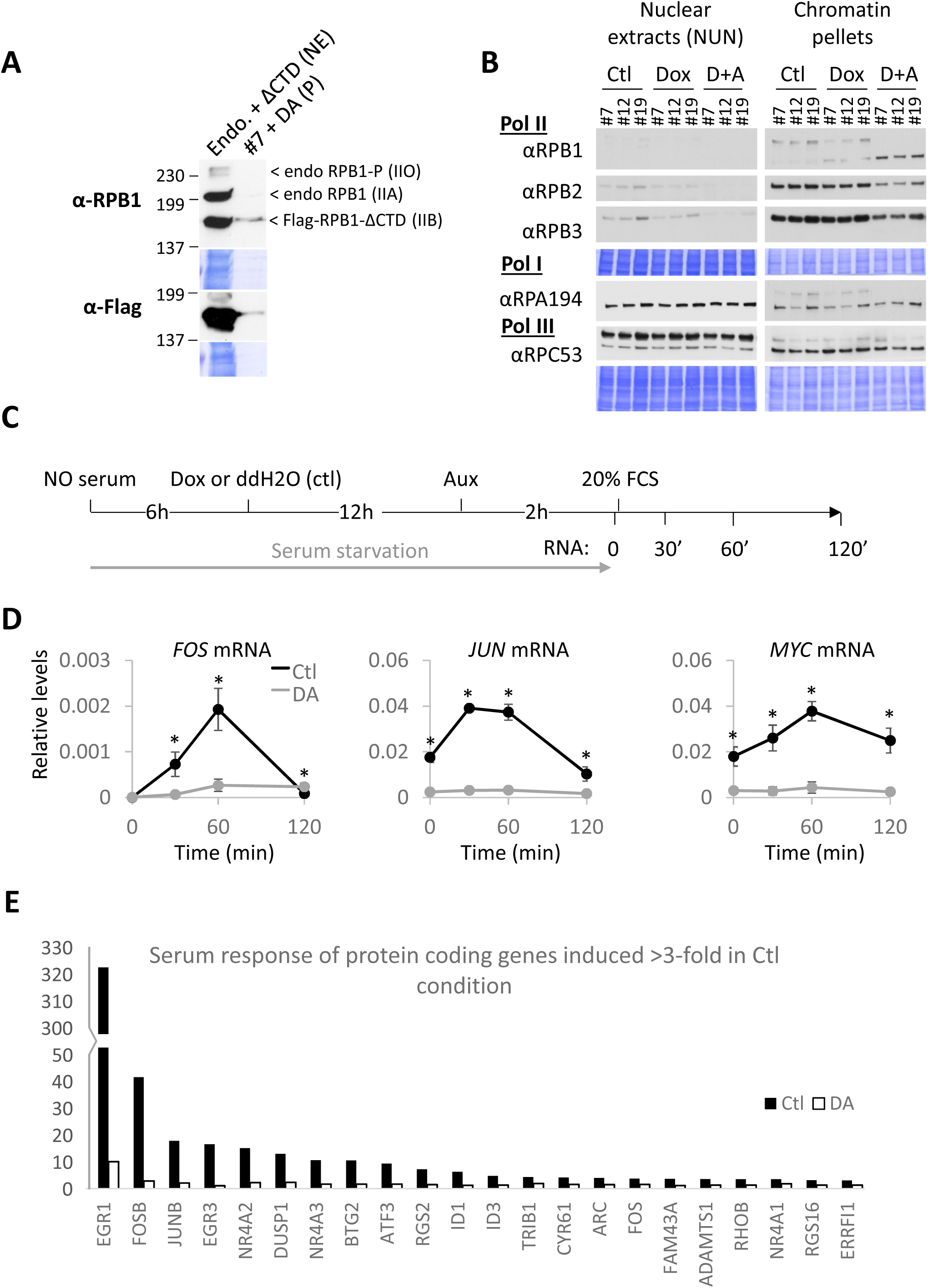
Related to Figure 2 and 5. **(A)** RPB1 and Flag immunoblots of nuclear extracts (NE) or chromatin pellets (P) prepared, respectively, from HeLa cells expressing endogenous RPB1 (Endo) and a Flag-tagged RPB1 lacking the entire CTD (ΔCTD) orDA-treated cells. The truncated RPB1 is present only on the chromatin-bound fraction in DA-treated cells. **(B)** Immunoblots and corresponding Coomassie-stained membranes of NUN extracts and pellets from untreated, DOX-treated or DA-treated cells probed for expression of different polymerase subunits. The physical isolation method used efficiently separates transcribing Pol III from Pol III containing phosphorylated RPC53 (slower migrating bands in nuclear extracts), presumably inactive (Lee et al., 2015). **(C)** Protocol used for time course analysis of RNA expression in serum-deprived cells following serum stimulation (also for Figure 6G-I). **(D)** Time course analysis of relative mRNA levels of selected immediate early genes in serum-starved RPB1-expressing (Ctl) and RPB1-depleted (DA) cells prior (Time 0) or after stimulation with serum as shown in **(C).** Data represent the average +/- SD. * indicates statistically significant differences (T-test; p<0.05). **(E)** Histogram of all protein-coding serum-inducible genes in Control (Ctl) and DA-treated cells revealed in a microarray analysis of pools of RNAs prepared from all three clones showing complete abolishment of the serum induction (the remaining enhancement observed in DA-treated cells corresponds to the fraction of cells that do not respond to the DA-treatment, see Figure 1F).

**Figure S3.**
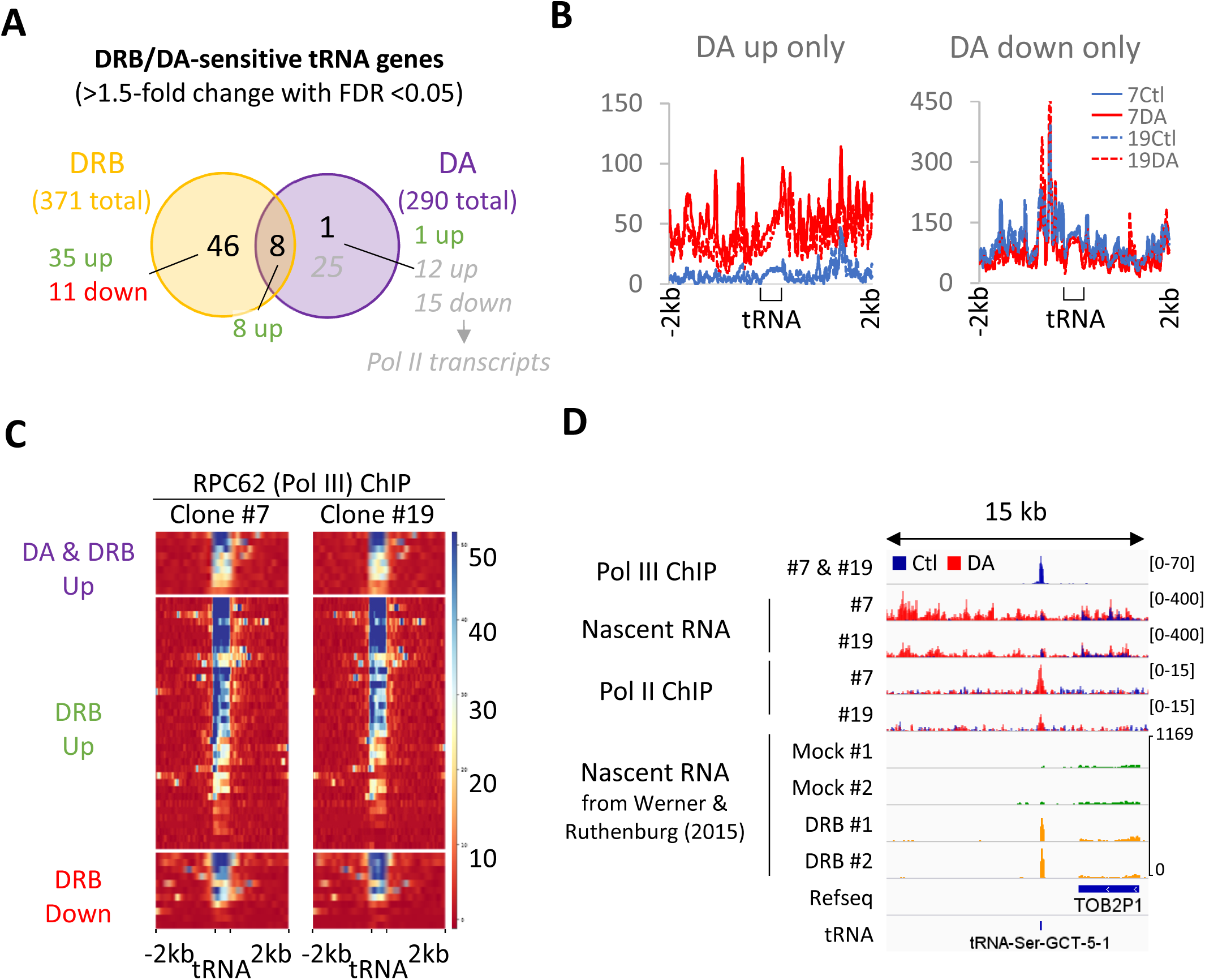
Related to Figure 3. **(A)** Venn diagram showing the number of tRNA genes affected >1.5-fold with a FDR <0.05 in DRB-treated wild-type HEK293 cells and DA-treated clones #7 and #19. Numbers in gray indicate numbers of tRNA genes affected by overlapping Pol II transcripts (and hence not considered in following analyses). **(B)** Nascent-RNA profiles of tRNA genes appearing up- or down-regulated because of changes in overlapping Pol II transcripts (CTD-less Pol II read-through or *bona fide* Pol II genes). Control conditions for clone #7 and #19 are shown in red, DA-treated profiles in blue. **(C)** Heat maps of RPC62 enrichment at tRNA loci in all 3 groups defined in Figure 4B, demonstrating that these loci are Pol III target genes in these two clones. **(B**) Example of a tRNA locus upregulated in DRB but not in DA-treated cells (second group in Figure 3C). The data correspond, in descending order, to input-subtracted RPC62 ChIP-seq (overlay of clone #7 and #19 data), overlays of normalized nascent RNA and input-subtracted RPB3 ChIP-seq coverage in RPB1-expressing (Ctl, blue) or RPB1-depleted (DA, red) clone #7 and #19 cells. In addition, nascent RNA coverage replicates of DRB- or mock-treated cells are shown, respectively, in green and yellow. Note that the presence of read-through transcription in DA-treated cells is accompanied by an accumulation of presumably transcribing CTD-less Pol II complexes at the Pol III locus.

**Figure S4.**
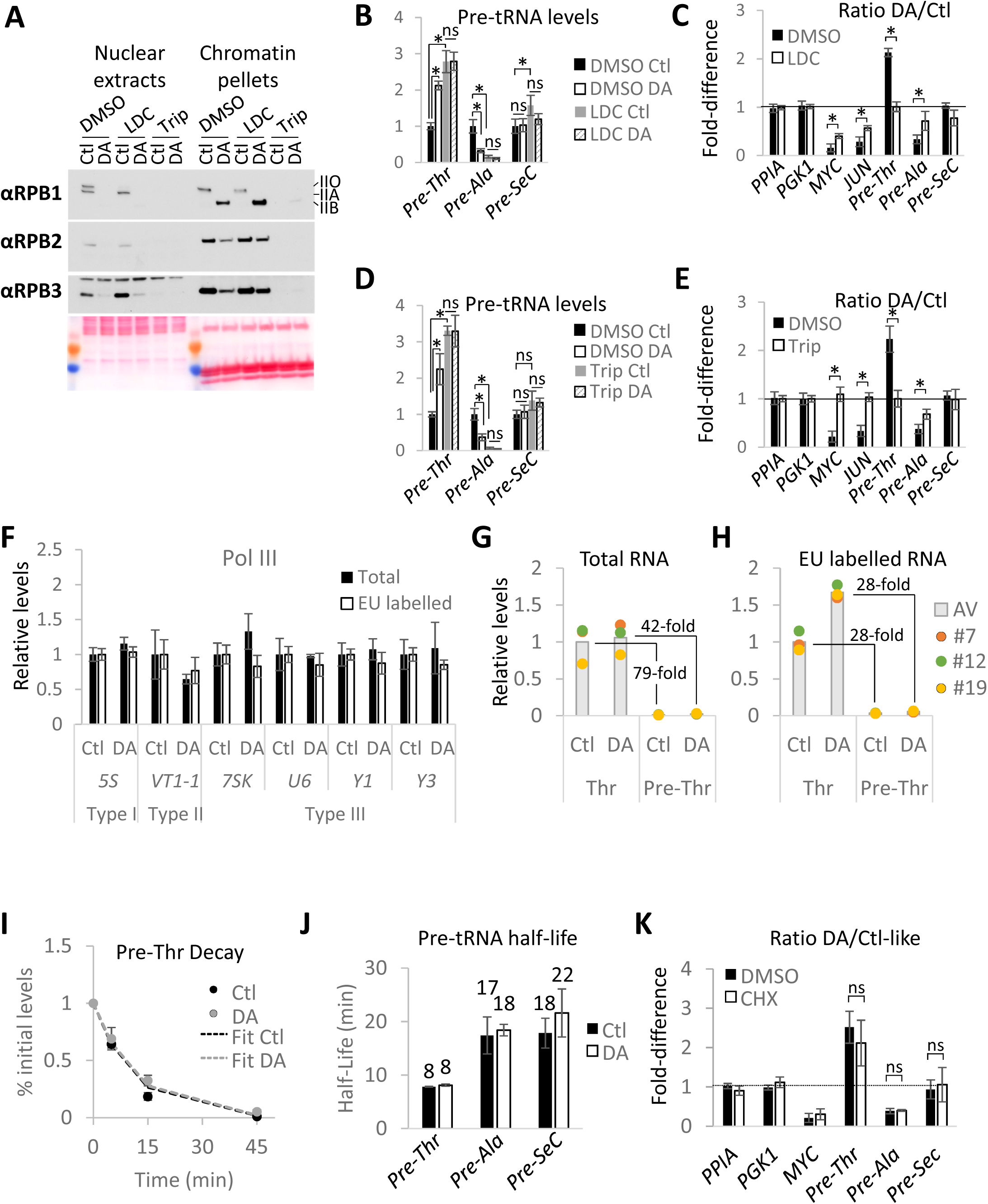
Related to Figure 4. **(A)** Immunoblots and corresponding Ponceau-stained membranes of NUN extracts and chromatin pellets prepared from RPB1-expressing (Ctl) or RPB1-depleted (DA) clone #7 cells pretreated with DMSO, LDC or Trip for 1 hr prior to auxin addition. IIO, IIA and IIB indicate the positions, respectively, of phosphorylated-, unphosphorylated- and truncated forms of RPB1. **(B)** Relative pre-tRNA expression levels in all three clones normalized to DMSO Ctl in RPB1-expressing (Ctl) or RPB1-depleted (DA) cells pre-treated with LDC or DMSO. **(C)** DA/Ctl ratios for data presented in **(B) as well as for some mRNAs. (D)** Same as **(B)** but with cells pre-treated with Triptolide and DMSO. **(E)** Same as **(C)** but for data shown in **(D). (F)** Relative expression levels of different types of Pol III-transcribed RNAs (other than tRNAs) in RPB1-expressing (Ctl) or RPB1-depleted (DA) cells for all 3 clones normalized to their levels in Ctl condition. Data shows levels in total (black bars) or metabolically labelled RNAs (white bars). **(G)** Relative levels of mature and precursor pre-Thr tRNA in total RNA prepared from RPB1-expressing (Ctl) and RPB1-depleted (DA) cells for all three clones. The histograms represent the average expression. **(G)** Same as **(H)** but in EU-labelled RNA. This shows that despite the fact that the primers used to measure the mature form also hybridize to the precursor form, the contribution of the labelled precursor is negligible (1/28 of the mature tRNA level). **(I)** Decay curve fits used to calculate the stability of pre-Thr tRNA in RPB1-expressing (Ctl) and RPB1-depleted (DA) cells for all three clones treated with ActD to block transcription. The data are shown as % of the levels in cells prior to ActD addition (Time 0). **(J)** Half-life of precursor tRNAs in RPB1-expressing (Ctl) or RPB1-depleted (DA) cells for all three clones. **(K)** DA/Ctl ratios of RNA expression levels in RPB1-expressing (Ctl) or RPB1-depleted (DA) cells for all 3 clones pre-treated 15 min with cycloheximide (CHX) or DMSO. Data in **(B)-(F)** and **(I)** to **(K)** represent the average +/- SD. * indicates statistically significant differences, ns non-significant differences (T-test; p<0.05).

**Figure S5.**
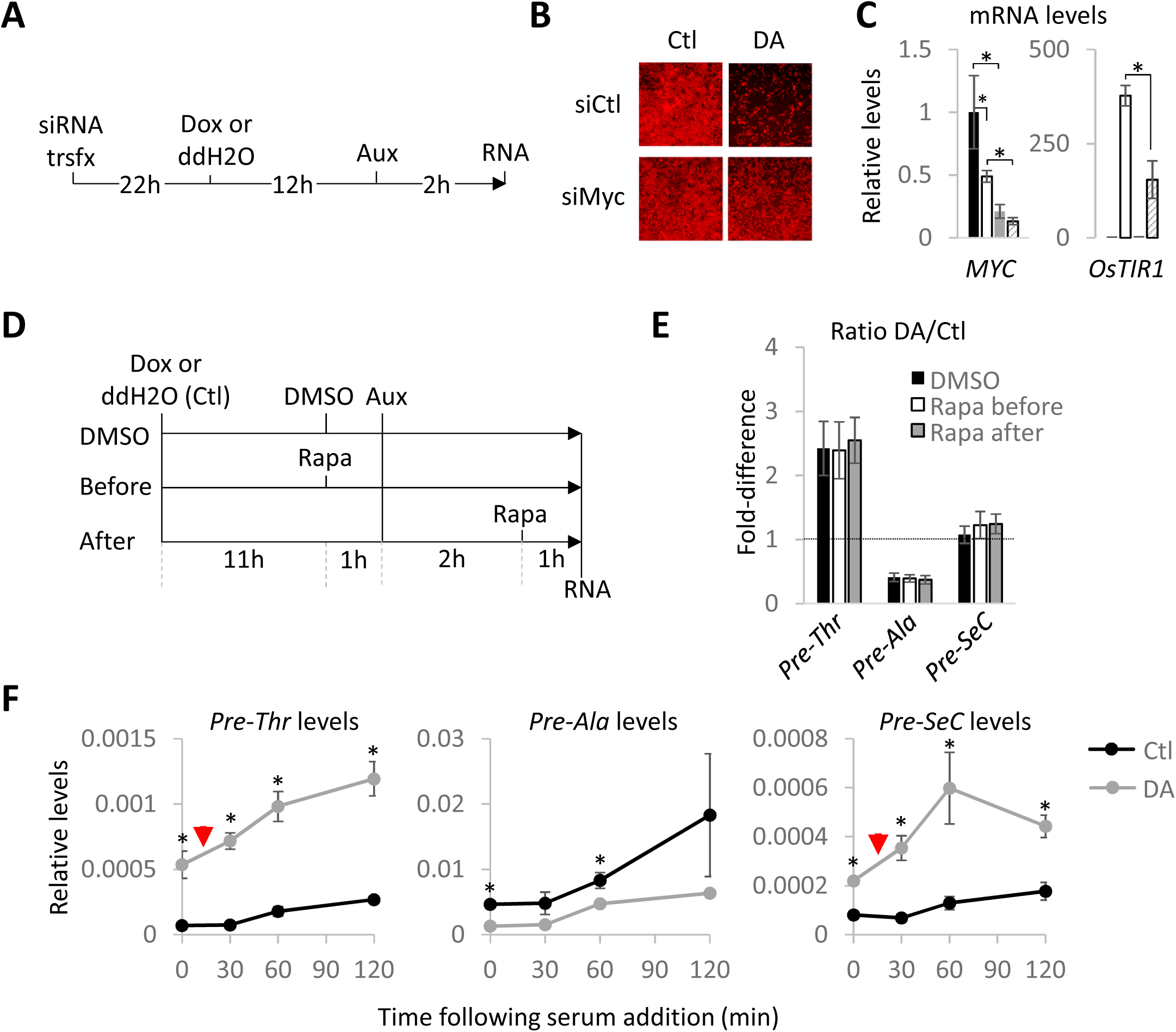
Related to Figure 5. **(A)** Protocol for siRNA-based knock-down prior to RPB1 depletion. **(B)** Fluorescence microscopy pictures showing the effects of MYC knock-down on RPB1 depletion in DA cells. **(C)** Relative mRNA expression levels normalized to siCtl-treated cells for all 3 clones transfected with a non-targeting (siCtl) or *MYC*-targeting pool of siRNAs (siMyc) prior to RPB1 depletion. **(D)** Protocol for rapamycin (Rapa) treatments prior to or after RPB1-depletion. **(E)** DA/Ctl ratios for data presented in (Figure 5F) for pre-tRNAs. **(F)** Time course analysis of relative pre-tRNAs levels in serum-depleted RPB1-expressing (Ctl) or RPB1-depleted (DA) cells stimulated with 20% FCS after RPB1 depletion. Red triangles emphasize the faster response of DA-treated cells during the first 30 min of serum stimulation. Data in **(C)-(F)** represent the average +/- SD. * indicates statistically significant differences (T-test; p<0.05).

**Figure S6.**
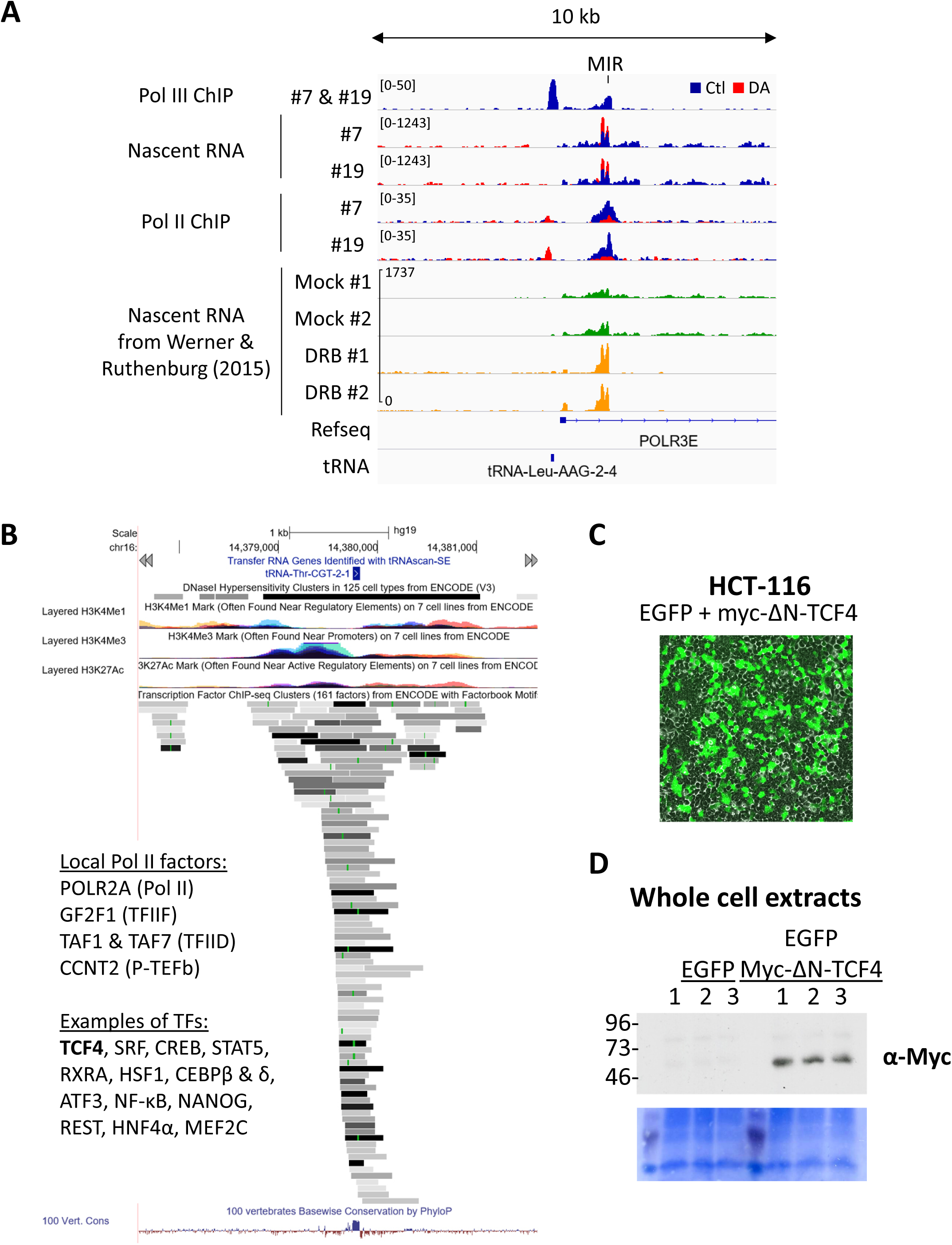
Related to Figure 6. **(A)** Input-subtracted RPC62 ChIP-seq (overlay of clone #7 and #19 data), overlays of normalized nascent RNA and input-subtracted RPB3 ChIP-seq coverage in RPB1-expressing (Ctl, blue) or RPB1-depleted (DA, red) clone #7 and #19 cells at a *MIR* locus that is transcribed by Pol III in the anti-sense direction from the Pol II-transcribed *POLR3E* gene (encoding a subunit of Pol III itself). In addition, nascent RNA coverage replicates of DRB- or mock-treated cells are shown, respectively, in green and yellow. The loss of elongating Pol II at this locus is correlated with an increase in transcription of the *MIR* element in both DA-treated cells and DRB-treated cells**. (B)** UCSC bowser snapshot of the tRNA-Thr-CGT-2-1 locus displaying ENCODE ChIP-seq data for common histone modifications found at promoters and regulatory regions as well as all available data for Pol II, its associated general transcription factors and the sequence-specific transcriptions factors immunoprecipitated in the vicinity of this locus, suggesting the presence of a local Pol II promoter near this tRNA locus. (C) Microscopy picture showing transfection efficiencies (EGFP-positive cells) of HCT116 cells co-transfected with plasmids and EGFP and a ΔN-TCF4 constructs. (D) Immunoblot and corresponding Coomassie-stain membrane of total extracts prepared form HCT116 cells transfected cells probed with anti-MYC antibody to evaluate expression of the myc-tagged ΔN-TCF4 construct.

**Figure S7.**
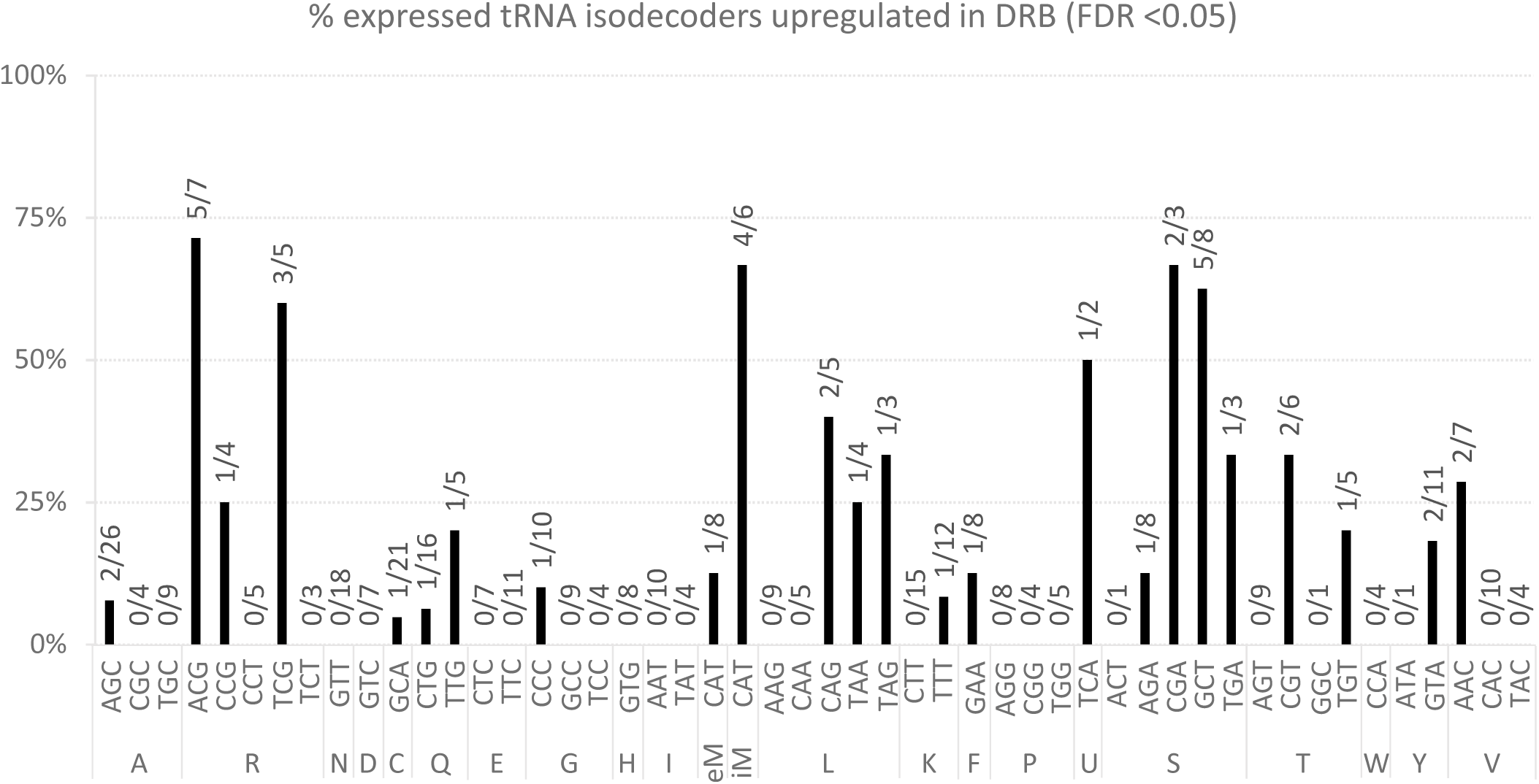
Related to Figure 7. Histogram showing the proportion (and numbers) of tRNA genes (grouped by anticodon) induced by DRB. See Discussion for details.

